# Climate History and Metabolic Trade-offs shape the Bacterial Response to Drought

**DOI:** 10.1101/2024.09.03.611070

**Authors:** Nicholas J. Bouskill, Stephany S. Chacon, Daniela F. Cusack, Lee H. Dietterich, Liang Chen, Aizah Khurram, Jana Voriskova, Hoi-Ying N. Holman

## Abstract

Soil drying challenges microbial viability and survival, with bacteria employing various mechanisms to respond to shifts in osmolarity, including dormancy or metabolic upregulation of osmoprotectants. However, the extent to which these responses are shaped by an organism’s phylogeny or the climate history of a given environment is poorly understood. This study examines the responses of phylogenetically similar bacteria from semi-arid and humid tropical forest soils to osmotic and matric stress using synchrotron radiation-based Fourier Transform Infrared spectromicroscopy. This non-destructive approach depicts the biochemical phenotype for whole cells under control and stress conditions. We observed that, under osmotic stress, bacteria upregulated cell-signaling pathways, rapidly turned over lipid-storage compounds, and increased osmolyte production. In contrast, matric stress induced a more muted response, typically elevating the production of carbohydrate stress compounds, such as glycine betaine and trehalose. While phylogenetically similar bacteria showed comparable biochemistry under control conditions, climate history played an important role in regulating responses to stress, whereby a stronger metabolic response was observed from semi-arid relative to tropical forest isolates. We conclude that bacterial stress response to drought can be more diverse than previously observed, and regulated by both phylogeny and climate history.

## Introduction

Environmental fluctuations - such as changes in pH, temperature, and moisture - can alter the chemical composition of the local environment^1–3^, and provoke both short-term (seconds to minutes) and long-term (hours to days) phenotypic and metabolic responses in microbial communities^4^. Soil microorganisms exhibit a range of adaptive traits to protect cellular integrity and maintain metabolism amidst these variable conditions^3,5^. Moreover, the frequency of environmental changes (occurring on hourly, diel, or seasonal scales), can engender anticipatory responses in individuals and microbial populations that selects for distinct life-history strategies^6–9^ and the emergence of seasonally recurring patterns^10^.

Soil drying represents a distinct physiological challenge to microbial communities. As soil dries, water forms thin films around soil particles, concentrating aqueous pore water constituents (dissolved nutrients, solutes, toxicants) while limiting the diffusion of substrates and extracellular enzymes^2,11^. This drying leads to osmotic stress from increasing solute and toxicant concentrations and matric stress from higher surface tension and capillary-bound water, both of which challenge the maintenance of turgor pressure, growth, and cellular viability^13,14^.

Microorganisms exhibit broadly different sensitivities to drought. Gram negative bacteria appear more sensitive to changing soil moisture compared to gram positive bacteria due to differences in cell wall structure^15^, which affects the regulation of intracellular osmotic pressure. To mitigate drought stress, bacteria can upregulate pathways to reduce intracellular osmotic potential^16,17^, primarily through the synthesis or accumulation of organic molecules (compatible solutes), like trehalose, and nitrogenous amino acids, such as ectoine, and glycine betaine^18^.

While some common responses of soil bacteria to drought have been described in the literature, a detailed understanding of specific bacterial mechanisms underpinning survival under rapid changes in osmotic potential is lacking. Microbial communities in ecosystems with significant environmental fluctuations are generally less sensitive to stochastic disturbances compared to those in more stable environments (see Hawkes and Keitt^22^ for greater discussion), and the legacy of drought and extent of soil drying within a given region can all influence the ecological niche of an organism, the traits expressed under stress, and the response to perturbation^6,20,21^. This raises the question of whether climate history and drought legacy drive microbial stress response. If so, microorganisms from the similar environments (e.g., humid tropical forest soils) should show consistent responses to osmotic stress. Conversely, if microbial responses to stress are hardwired through the phylogenetic conservation of functional traits^19^, then phylogenetically-related organisms should show similar responses to stress regardless of environment.

We address these questions by analyzing the metabolic responses of four common soil bacteria - two gram-positive, *Arthrobacter pascens* and *Bacillus thuringiensis,* and two gram-negative bacteria, *Enterobacter asburiae* and *Sphingomonas paucimobilis -* to stress conditions typical of drought (Fig. 1, Supplementary table 1). Phylogenetically similar representatives of each of the four phyla (Fig. 2a) were isolated from two different geographic environments; a semi-arid montane (termed SAM isolates) inceptisol, and an ultisol within a humid tropical forest (termed HTF isolates).

**Figure 1:**
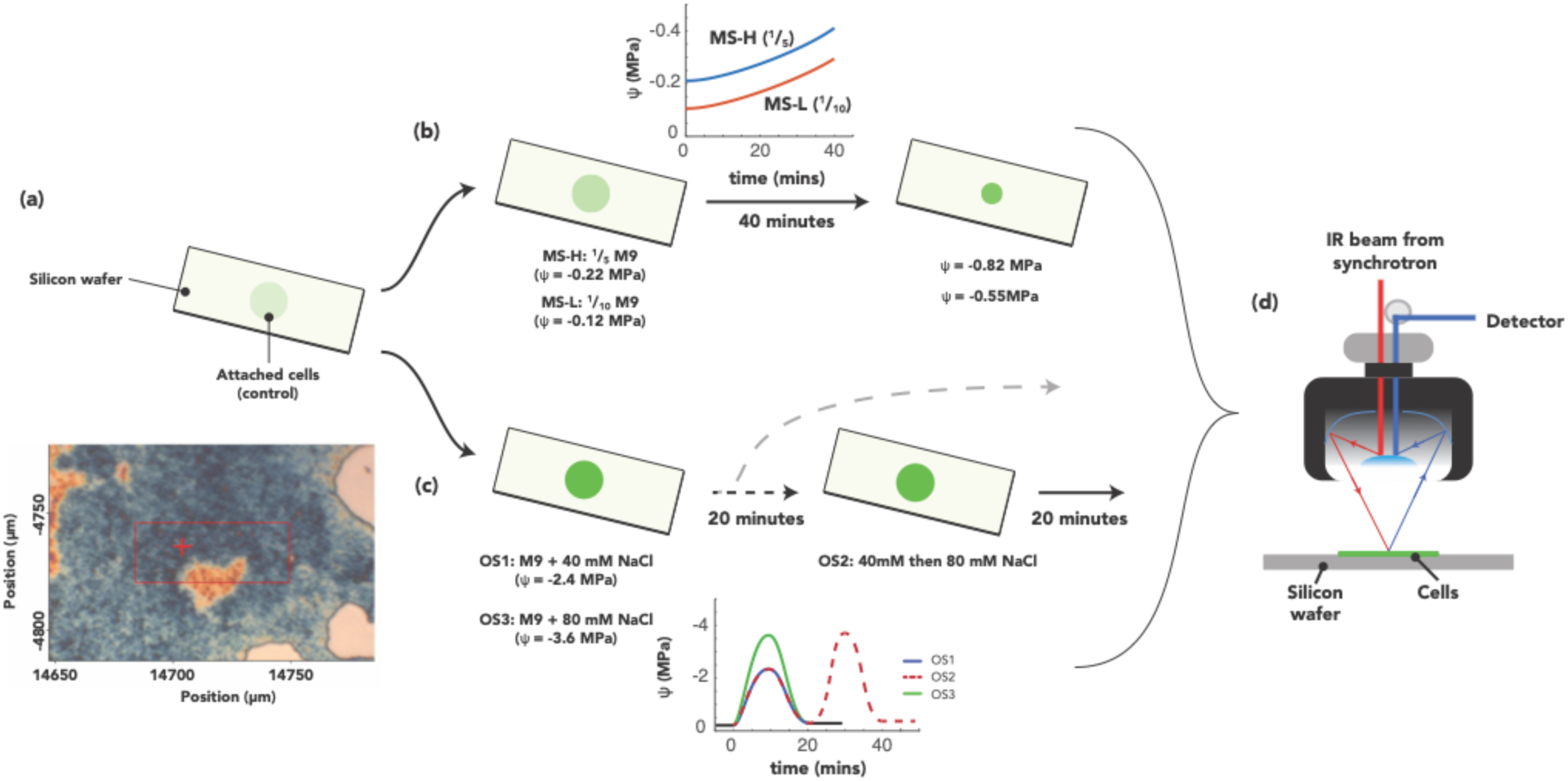
A schematic of the experimental approaches manipulating the water potential for synchrotron-based infrared spectromicroscopy measurements. Details are provided in the materials and methods, however, briefly, panel (a) depicts cell preparation prior to exposure, whereby, a 4 μl aliquot of an exponentially growing axenic cell culture is allowed to adhere to a silicon wafer to form microcolonies (insert), which are either exposed to (b) two different matric stress experiments (MS), where M9 media of different dilution strengths are allowed to evaporate and concentrate the media. Media was either ⅕ strength of initial M9 media (MS-H) or 1/10 strength (MS-L). Isolates were also exposed to (c) three distinct osmotic stress experiments. Two osmotic stress experiments briefly pulse the media with NaCl at a low (40 mM = OS1) and either imaged cell biochemistry afterwards, or immediately pulsed the isolates with the high concentration of NaCl (i.e., a sequential step-up experiment of 40 mM to 80 mM = OS2). A final experiment exposed cells to the high (80 mM = OS3) concentration^21^. The phenotypic and metabolic response of the cells to stress is measured using (d) synchrotron-based FTIR, and analyzed through principal-components, linear-discriminant analysis (PC-LDA).

**Figure 2:**
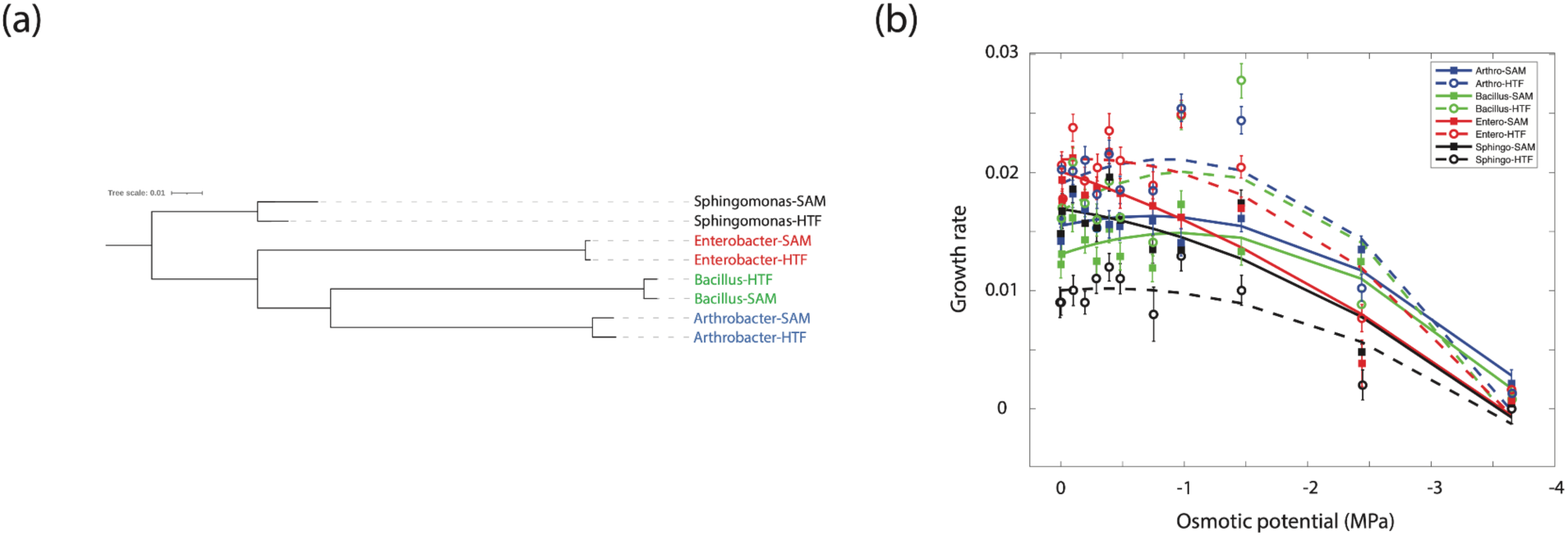
Phylogeny and growth of the different isolates. (a) Phylogenetic relationships between the various isolates. (b) Growth rates across a range of solute potential measurements. The rates are shown as a second-order polynomial that best fits the data. Phylogenetically similar bacteria share the same line color, whereas isolates from the HTF are depicted with dashed lines.

The semi-arid montane site (East River, Colorado, 38.92N, 106.9487W) experiences strong fluctuations in soil moisture, where snowmelt saturated soils give way to summer and fall drought, and osmotic potential can reach as low as -20 MPa prior to monsoon season. The humid tropical forest on Buena-Vista Peninsula, Panama (9.185N, 79.8266W), experiences shorter dry periods than the semi-arid site (∼ 133 days on average), with a more moderate decline in soil moisture. The comparison of isolates from these two ecosystems affords an examination of how phylogeny and differential climate history shape bacterial response to perturbation.

We categorize drought stress into the constituents of matric and osmotic stress (Fig. 1), each with mild and severe treatments, which may impose diverse metabolic costs on bacterial cells. The osmotic stress (OS) experiment, involves rapid changes in ionic potential, and also examines whether pre-conditioning cells, through exposure to a lower stress concentration, enables an acclimatory metabolic response as noted previously for bacterial populations^21^.

Examining physiological responses to stress necessitates the use of a sensitive and non-invasive measurement that has little to no impact on the cellular components (e.g., water, lipids, proteins, and carbohydrates, which comprise the basic structure of all bacterial systems). Infrared (IR) absorption spectroscopy in the mid-infrared region (4000 – 650 cm^-1^ in units of wavenumber or reciprocal centimeters) exploits the energy transfer from the infrared photons of specific vibrational frequencies to the distinct vibrational modes of the covalent bonds present in virtually all organic molecules. A highly beneficial feature of this non-invasive method is that these absorptions only occur at resonant frequencies, providing immediate information about the structure and composition of molecules in a biological sample from its mid-IR absorption spectrum even without *a priori* knowledge.

Here we used ultra-sensitive synchrotron radiation-based Fourier Transform Infrared (SR-FTIR) spectromicroscopy to measure changes in cellular composition and molecular structure originating from exposure to disturbance. This SR-FTIR approach can provide a 1000-fold increase in signal-to-noise (relative to an FTIR instrument using a halogen light source), for tracking dynamic cellular events in a 3- to 10-μm diameter area. It has previously tracked exposure events over time in a high-throughput and multiplexed manner and has been used to characterize changes in this biochemical composition within fluctuating environments^23–25^.

Herein we describe microcolony experiments aimed at testing the following hypotheses. (1) Short-term osmotic and matric stress provoke biochemical responses distinct from the control biochemistry regardless of an organism’s phylogeny or climate history. (2) Climate history will shape the metabolic response to stress, whereby organisms adapted to seasonal drought will show a stronger metabolic response (through elevated osmolyte production) than isolates from humid tropical forest soils.

## Results

### Growth rates and survivorship measurements emphasize the non-lethal nature of the experimental stress

To explore whether broad differences in cellular survival and growth rate under drought stress were apparent between bacteria isolated from the SAM and HTF soils, we first measured survival of cells undergoing the matric stress (MS) and osmotic stress (OS) experimental conditions using flow cytometry. Under these conditions no significant differences were noted in survivorship between the control and experimental treatments, with > 97 % of cells alive following exposure to either of the stress conditions (Supplementary Fig. 1), demonstrating the non-lethal nature of the current experiments. Next, we measured the growth rate of the different isolates under increasing concentrations of NaCl (between 2 - 750 mM, spanning an added solute potential gradient of ∼0 to -3.6 MPa) as a proxy for drought stress. We observed climate history to play a role in separating the growth rate of isolates. The HTF isolates, with the exception of the Sphingomonas-HTF, had the highest growth rate, and showed greater sensitivity to increasing solute potential relative to SAM isolates. In general, increasing solute concentration imparted a threshold effect on growth rate, whereby osmotic potential greater than -1 MPa resulted in declining growth rates (Fig. 2b). However, the gram-positive isolates from the SAM soils had the highest growth rates at higher solute concentrations, and a gentler decline in growth rate overall.

### Multivariate infrared spectroscopy illuminates the bacterial biochemical phenotype

Infrared photons from the broad-band synchrotron light source were focused through an IR microscope onto a thin layer of bacteria under control (i.e., no stress), osmotic stress, or matric stress conditions. The view-field of the bacterial layer was divided into equal-sized 5-µm×5-µm squares before raster scanning, collecting full SR-FTIR spectra at each pixel. Each spectrum has 8,192 dimensions (variables), which represent signals from fatty acid esters, lipids, proteins, and carbohydrate absorptions. We focused our analysis on the spectral features in the cellular-biochemical region (1,800 - 1000 cm^-1^), reducing the dimensions of each spectrum from 8,192 to 1,739 variables, which still capture infrared signals from functional groups of lipids, proteins, and carbohydrates (Supplementary Fig. 2a). We first treated the dataset with mean-centered principal component analysis (PCA), as a purely exploratory and unsupervised way to extract the important factors accounting for the dataset’s maximum variance.

The PCA is highly influenced by outliers in our dataset. To better distinguish between isolates and their responses to different treatments, we used the first five PC scores in a linear discriminant analysis (LDA) algorithm. Principal Component-Linear Discriminant Analysis (PC-LDA) allowed us to compared the infrared spectral features of the eight bacteria isolates and identify the chemical bonds responsible for cluster separation (Supplementary Fig. 2c). Results are visualized via score plots (Fig. 3), which display each spectra as a point in multidimensional space, and through cluster vector spectra (Supplementary Fig. 3a), that are reconstructed mean spectra inside a vector space following PC-LDA data reduction. Each cluster vector spectrum uses the global mean cluster spectrum as the baseline and all individuals are quantitatively compared to it.

**Figure 3:**
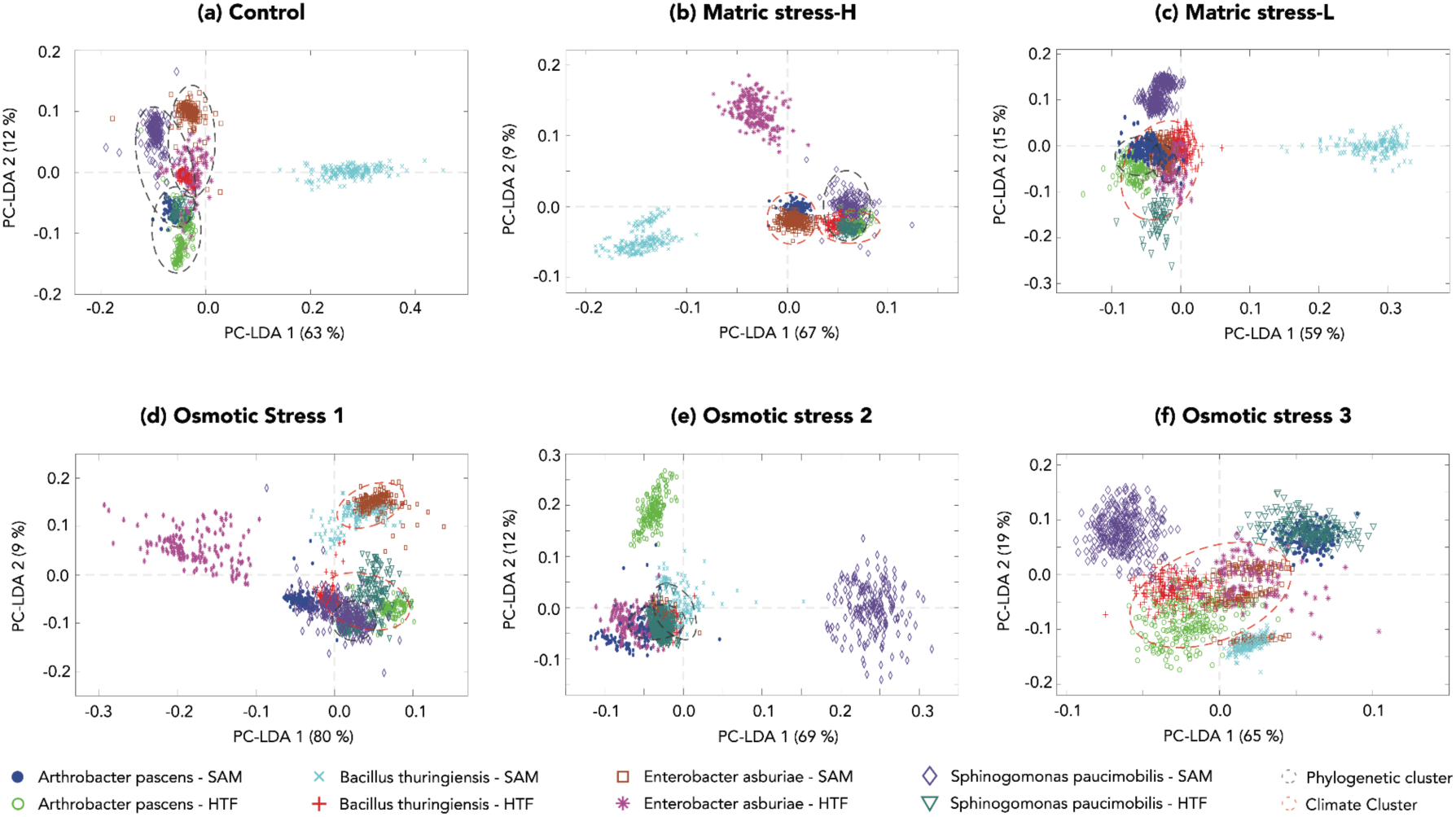
Ordination plots based on Principal Component-Linear Discrimination Analysis derived from infrared spectral biochemical fingerprints of multiple cells within (a) the control experiments, and (b-f) the different stress experiments. Each point on the ordination represents several cells within a 5-10 micron diameter spot area.

### Diversity in biochemical phenotypes under stress-free control conditions

Bacterial microcolonies without induced stress generally cluster by phylogeny, with separation by cell wall structure and intracellular energy storage polymers (Fig. 3a, Supplementary Fig. 3b). The Gram-positive Bacillus from semi-arid soils (Bacillus-SAM) exhibits distinct biochemistry relative to all other isolates along the primary axis (PC-LDA1), which accounted for ∼64 % of the variance. This axis is primarily defined by spectral features affiliated with bacterial energy storage polymer mixtures, such as polyhydroxyalkanotes (PHA), polyhydroxybutyrates (PHB), and energy yielding phospholipids (Supplementary Fig. 3a panel f). The largest contribution to the separation lies in the lipid/fatty acid hydrogen-bonded >C=O region 1750-1710 cm^-1^, which consisted of several overlapping peaks (Supplementary Fig. 2c panel f) at ∼1745 cm^-1^, ∼1740 cm^-1^, and 1732 cm^-1^ assigned to υ(>C=O) of the ester within PHB or within PHA and PHB mixtures^26,27^. The largest bands at ∼1730 cm^-1^ were assigned to υ(>C=O) of the carboxylic acid (–COOH) of phospholipids with additional supporting contributions from protein amide II (∼1542 cm^-1^, 1518 cm^-1^), δ_as_ and δ_s_ (C ̶ H) of CH_3_ and CH_2_ (∼1453 cm^-1^, ∼1338 cm^-1^), υC-O of ester (∼1290 cm^-1^), υ_as_ (>PO_2_^−^) at ∼1240-1220 cm^-1^ and υ_s_ (>PO_2_^−^) at ∼1090 - 1080 cm^-1^ of phospholipids and nucleic acids, and overlapping υ (C-O-C) and υ(P-O-C) at 1170 - 1160 cm^-1^ of carbohydrates^28–30^.

The secondary axis (PC-LDA2), which accounted for ∼13% of the total variance, was largely defined by absorbance of molecular features associated with cell wall structure (Supplementary Fig. 3a). Comparison of the secondary derivative spectra (Supplementary Fig. 2a) with the PC-LDA2 loading spectra (Supplementary Fig. 3b) show a number of spectral bands from the Gram-positive bacteria that can be assigned to the functional groups of bacterial cell-wall component peptidoglycan. The bands at ∼1725-1705 cm^-1^ are assigned to hydrogen-bonded carbonyl groups υ (>C=O) of aldehydes, ketone and carboxylic acids, while those at ∼1600 cm^-1^ and ∼1400 cm^-1^ relate to υ_as_ and υ_s_ (>C=O) of carboxylate (– COO ^−^) end or side chains of peptidoglycan. The bands at ∼1659 cm^-1^ and ∼1542 cm^-1^ are amide I and amide II of proteins arising in part from amino acids, whereas bands at ∼1455 cm^-1^ and ∼1380 cm^-1^ are δ_as_ and δ_s_ (C ̶ H) of CH_3_ and CH_2_, respectively. The bands at ∼1223 cm^-1^ and ∼1075 cm^-1^ are υ_as_ and υ_s_ (>PO_2_^−^) of phospholipids and nucleic acids. Finally, bands of the complex sugar aromatic ring vibration modes are at ∼1150, ∼1060 cm^-1^, and 1045 cm^-1^ ^29,31,32^.

Multivariate discriminative analysis showed bacteria from the same phylogeny but different climate ecosystems share similar biochemical composition. The cluster vector spectra (Supplementary Fig. 3c) for the two Arthrobacter isolates, for example, showed overlapping biochemical phenotypes under stress-free conditions, as shown by the contemporaneous peaks in the 1800-1000 cm^-1^ region, albeit at different peak intensities. This was also the case for the two Sphinogomonas and Enterobacter isolates (Supplementary Fig. 3c). The greatest distance between paired isolates was observed for the Bacillus, with infrared absorbance from Bacillus-SAM showing a distinct abundance in energy storage polymers relative to Bacillus-HTF.

### Diversity in biochemical phenotype under stress conditions

This pattern of separation of isolates based on cell-wall structure morphology and the energy storage polymers does not always hold under the different stress treatments (Fig. 3b-f). There is a degree of heterogeneity in response to stress, unrelated to cell-wall structure. Under several conditions, including MS-H, OS1, and OS3, organisms of different phylogeny isolated from sites with the same climate history (i.e., SAM or HTF) show a more similar response to stress, clustering together on the ordination. Conversely, there is also evidence for a similar response based on phylogeny rather than climate history for the MS-L and OS2 experiments. A detailed account of the factors separating different isolates under stress is provided in the supplementary material.

### Changes to the biochemical phenotype under stress

Figure 4 illustrates the biochemical phenotypic properties of cells and the overall dissimilarity between control cells and those under OS or MS stress conditions. The majority of variance between the various experiments is captured by the first two factors of the PC-LDA analysis, explaining over 80% of the variance (with the first 3 PC-LDA components explaining over >90 %, Supplementary Fig. 4a,b), allowing us to focus on inter-treatment variation for all eight isolates.

**Figure 4:**
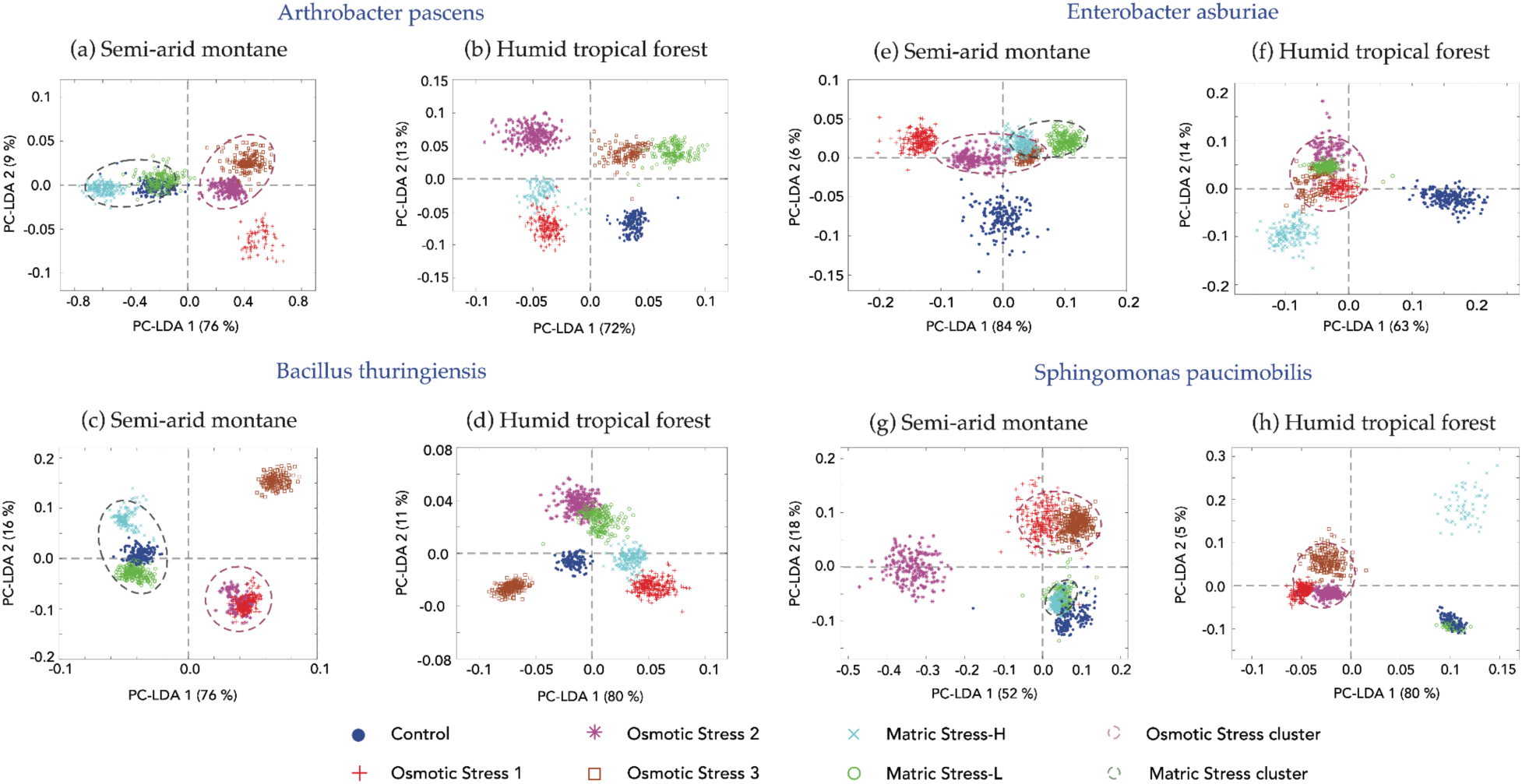
Ordination plots based on Principal Component-Linear Discrimination Analysis for each isolate. Each panel shows the ordination of the biochemical phenotype under control and different experimental conditions of osmotic and matric stress. The panels provide results for bacteria isolated from semi-arid montane (SAM) and humid tropical forest (HTF) soils for the gram-positive *Arthrobacter pascens* (a/b), and *Bacillus thuringiensis* (c/d), and the gram-negative *Enterobacter asburiae* (e/f), and *Sphingomonas paucimobilis* (g/h). The dashed ovals cluster similar experimental responses to either osmotic stress (brown dashed lines) or matric stress (green dashed lines).

To highlight the broad responses of different isolates to stress, we analyzed spectra from specific regions: the lipid/fatty acid region (∼1760 - 1600 cm^-1^), the phosphorus bond region (1230 - 1200 cm^-1^), and the carbohydrate region (1250 - 1000 cm^-1^) (Fig. 5 & 6). The phosphorus bond region can be indicative of the presence or absence of energy storage polymers, and molecules involved in two-component signal transduction systems, which consist of >PO_2_^−^ groups. We also contextualize these broad responses with data on shifts in microbial storage compounds (e.g., PHAs, PHBs, polyphosphates, and glycogen), and stress related metabolites (e.g., trehalose, glutamate and proline) and osmolytes (e.g., mannitol, ecotoine, and glycine betaine), identified by their characteristic IR spectra (Supplementary Fig. 2b and Supplementary Table 3). Detailed analysis of band position shifts relative to the control spectra is provided in the Supplementary Tables 4 to 11.

**Figure 5:**
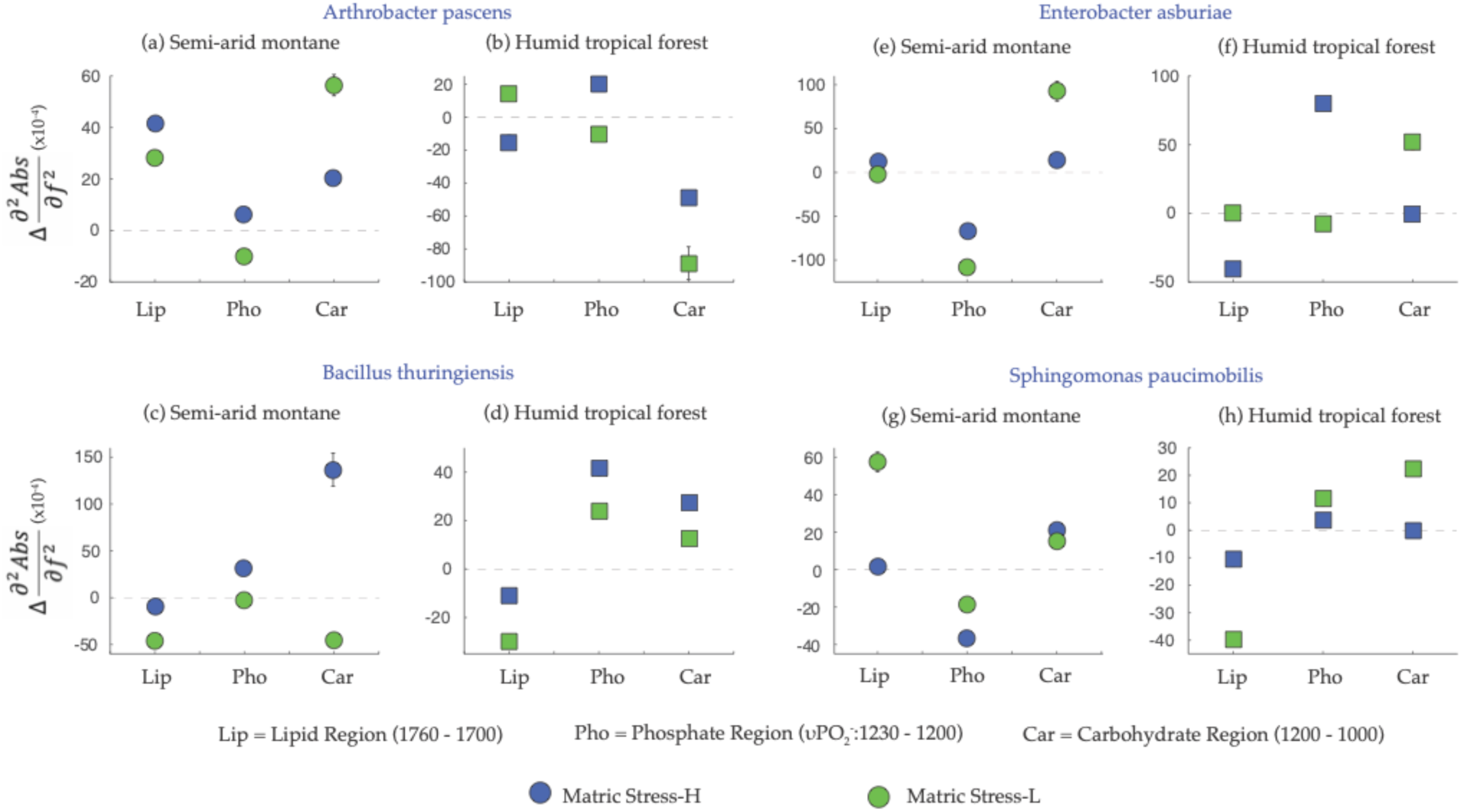
Metabolite responses to the matric stress experiments for each isolate. The figure depicts the aggregate shifts in the regions representing lipids and fatty acids (1760 - 1700 cm^-1^), the phosphate region (1250 - 1200 cm^-1^), and the broader carbohydrate region (1200 - 1000 cm^-1^) as shown by the mean relative changes (+/- 10% errors) of the amplitude of secondary derivative absorption peak spectra. These shifts are normalized by the control spectra.

The short-pulsed nature of the stress experiments was capable of provoking a metabolic response in each of the isolates, however, the divergence in biochemical phenotype between the treatments and control was dependent on the climate history of an isolate (Fig. 4). For example, the biochemical phenotype of experimentally stressed isolates from tropical forest soils never overlapped with the control phenotype (Fig. 4b, d, f, h), even under the mild matric stress experiments. This is demonstrated by Gram-positive Arthrobacter-HTF and Bacillus-HTF, and the Gram-negative Enterobacter-HTF, where a unique biochemical phenotype was observed for the control and each of the stress conditions (Fig.4b). For Arthrobacter-HTF the first two PC-LDA components separated the matric stress (MS-H, MS-L) from osmotic stress experiments (OS1, OS2, OS3), as well as high stress (MS-H, OS2, OS3) from low stress experiments (MS-L, OS1). Bacillus-HTF showed clustering of OS and MS experiments (e.g., MS-H and OS1), with OS2 diverging along PC-LDA1. The biochemical phenotype of Enterobacter-HTF clustered for the OS experiments, with MS-H ordinating away from the other experiments along PC-LDA1.

For Sphingomonas-HTF (Fig. 4h), the matric and osmotic stresses were separated along the primary PC-LDA, with a strong cluster of OS experiments, however, there was overlap between the control experiments and MS-L.

Unlike the HTF isolates, those from semi-arid soils showed strong overlap between the biochemical phenotypes of matric stress experiments and the controls (Fig.4a, c, e, g). The Arthrobacter and Bacillus-SAM isolates showed co-clustering of the matric stress and control biochemistry, and separation from the osmotic stresses along the primary axis PC-LDA1 (Fig. 4a, c), which accounted for a majority of variance (76%). For the Gram-negative Sphingomonas-SAM, matric stress and control biochemistry co-clustered and were separated from osmotic stresses along the secondary PC-LDA axis (Fig. 4g). Finally, the Enterobacter-SAM control phenotype exhibited minimal overlap between the various stress experiments (Fig. 4e), similar to that of the Enterobacter-HTF isolate.

### Quantitative analysis of isolate-specific response to different stress

The biochemical response of the SAM isolates to MS-H stress showed an overall increase in the carbohydrate (1200 – 1000 cm^-1^) compared to the controls (Fig. 5). This increase is linked to higher osmolyte production (Fig. 7) and an accumulation of carbohydrate storage compounds like glycogen (Fig. 8). While the lipid region showed variable response, there was a general increase in the PHAs under stress (Fig. 8). In contrast, HTF isolates showed an overall upregulation of the phosphate (υPO_2_^-^) region (1230 - 1200 cm^-1^), and a general decline in the lipid and carbohydrate regions (Fig. 5). This broader decline in lipids was accompanied by a strong decrease in PHA storage compounds. Glycogen also decreased for Bacillus and Sphingomonas, however, there was a general increase in some osmolytes, mainly glycine betaine, but also trehalose and ectoine (Fig. 7).

Under the mild MS experiment, MS-L, isolates from most semi-arid soils showed consistent overlap between the emergent biochemical response under matric stress and the control biochemical phenotype (Fig. 4a,c,g). This emphasizes that MS-L did not impart significant stress on these isolates. Although there was a broad increase in carbohydrate production (Fig. 5), the osmolyte response was generally weak, with only some evidence of trehalose production in Enterobacter and Sphingomonas isolates (Fig. 7). In contrast, isolates from humid tropical forest soils displayed a more divergent response under MS-L compared to control cells, characterized by increased production of carbohydrates and a decline in lipids (Fig. 5). Specifically, PHA storage declined in most isolates as did most osmolytes (Fig. 7). The exception here was Sphingomonas-HTF, which increased the production of PHA relative to the control, and increased concentrations of glycogen, trehalose, and ectoine.

Several isolates, particularly from semi-arid soils (e.g., Arthobacter-SAM, Bacillus-SAM, and Sphingomonas-SAM) as well as Sphingomonas-HTF, showed distinct biochemical responses to osmotic stress compared to matric stress. Osmotic stress experiments produced unique biochemical responses across the SAM isolates relative to controls (Fig. 4a,c,g,h), with a broad decrease in the lipid region, and increase in the carbohydrate region. Under osmotic stress 1 (40 mM), the decrease in the lipid region cannot be explained by changes in storage compounds, as we observed a general stress-induced increase in PHA, PHB, and polyphosphate across all isolates (Fig. 8). The broader increase in carbohydrate was mirrored by a general rise in osmolytes of multiple types (Fig. 7). In contrast, the majority of HTF isolates (Arthrobacter-HTF, Bacillus-HTF, and Sphingomonas-HTF), showed a broad decrease in lipid and carbohydrate regions, with a concomitant increase in the υPO_2_^-^ region (Fig. 6). Despite the broader drop in carbohydrate, there was a strong increase in stress-related osmolytes, notably trehalose, and glycine betaine.

**Figure 6:**
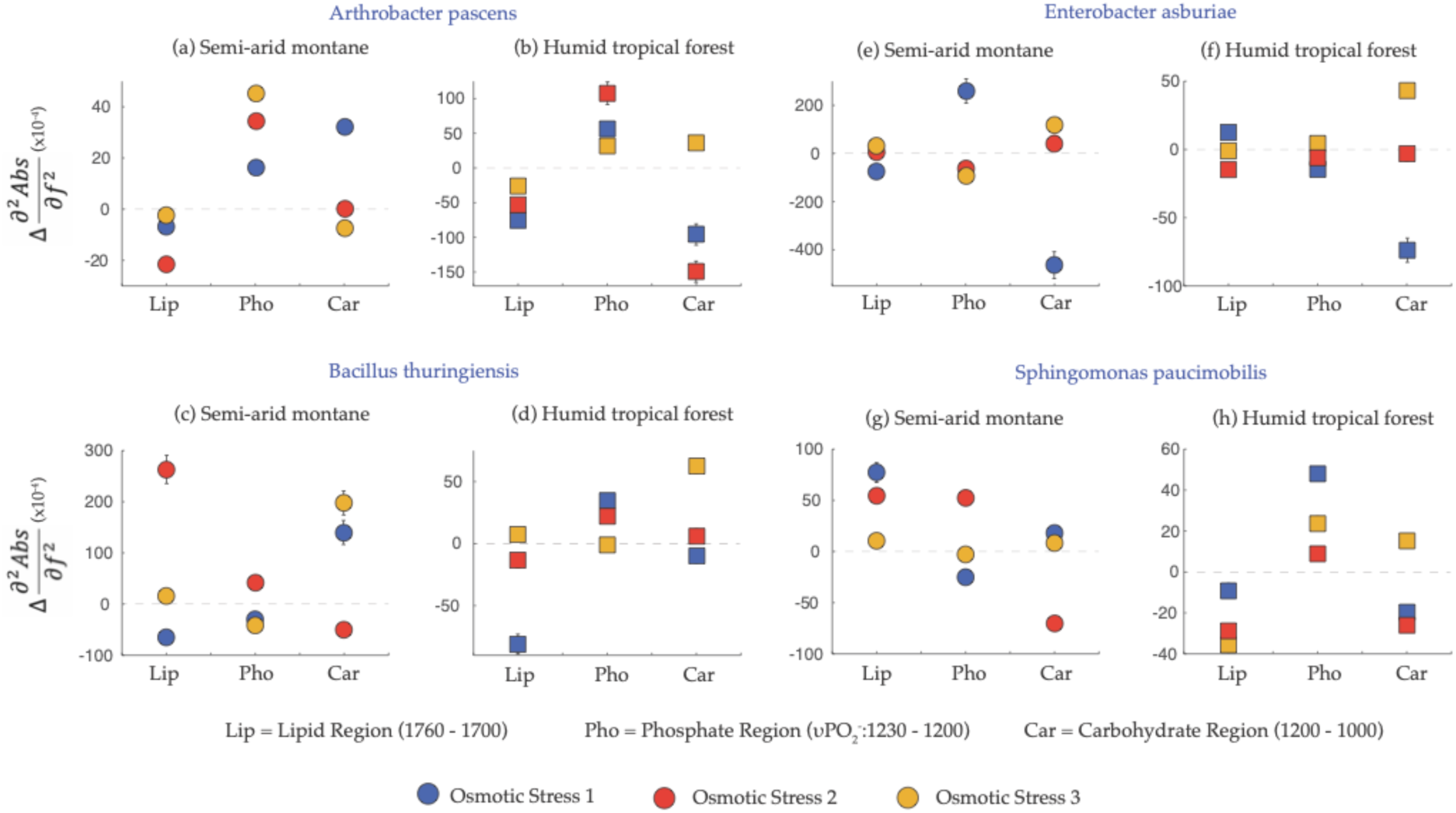
Metabolite responses to the osmotic stress experiments for each isolate. The figure depicts the aggregate shifts in the regions representing lipids and fatty acids (1760 - 1700 cm^-1^), the phosphate region (1250 - 1200 cm^-1^), and the broader carbohydrate region (1200 - 1000 cm^-1^) as shown by the mean relative changes (+/- 10% errors) of the amplitude of secondary derivative absorption peak spectra. These shifts are normalized by the control spectra.

Preconditioning the cells to lower osmotic stress before subjecting them to higher stress (OS2) prompted two major responses, (a) an intermediate biochemical response clustering between the OS1 and OS3 phenotypes (Fig. 4a,e,h), and (b) several unique responses in which the OS2 experiment clusters away from the other OS experiments (Fig. 4b,d,f,g). For the SAM isolates OS2 resulted in a decline or no change in the carbohydrate regions, distinct from the OS1 response. (Fig. 6). The lipid region response varied among isolates. In general, there was a similarly strong osmolyte response under OS2 as that observed in OS1, with glycine betaine a particularly prominent compound increasing under stress. Similarly, the storage compounds generally increased under the OS2 treatment, with PHA increasing within most isolates. In instances where PHA declined (e.g., Bacillus-SAM) there was a concomitant increase in glycogen.

For the HTF isolates, including Arthrobacter-HTF, Enterobacter-HTF and Sphingomonas-HTF, we observed a similar decrease in the carbohydrate region during OS2 as seen in the OS1 experiment (Fig. 6). This was accompanied by an increase in osmolyte production, notably glycine betaine (Fig. 7). In contrast, Bacillus-HTF showed an increase in the carbohydrate region, particularly in glycine betaine, under the step-up experiment. For most HTF isolates, this strong increase in osmolytes was accompanied by a decline in cellular storage compounds, including glycogen and PHA.

**Figure 7:**
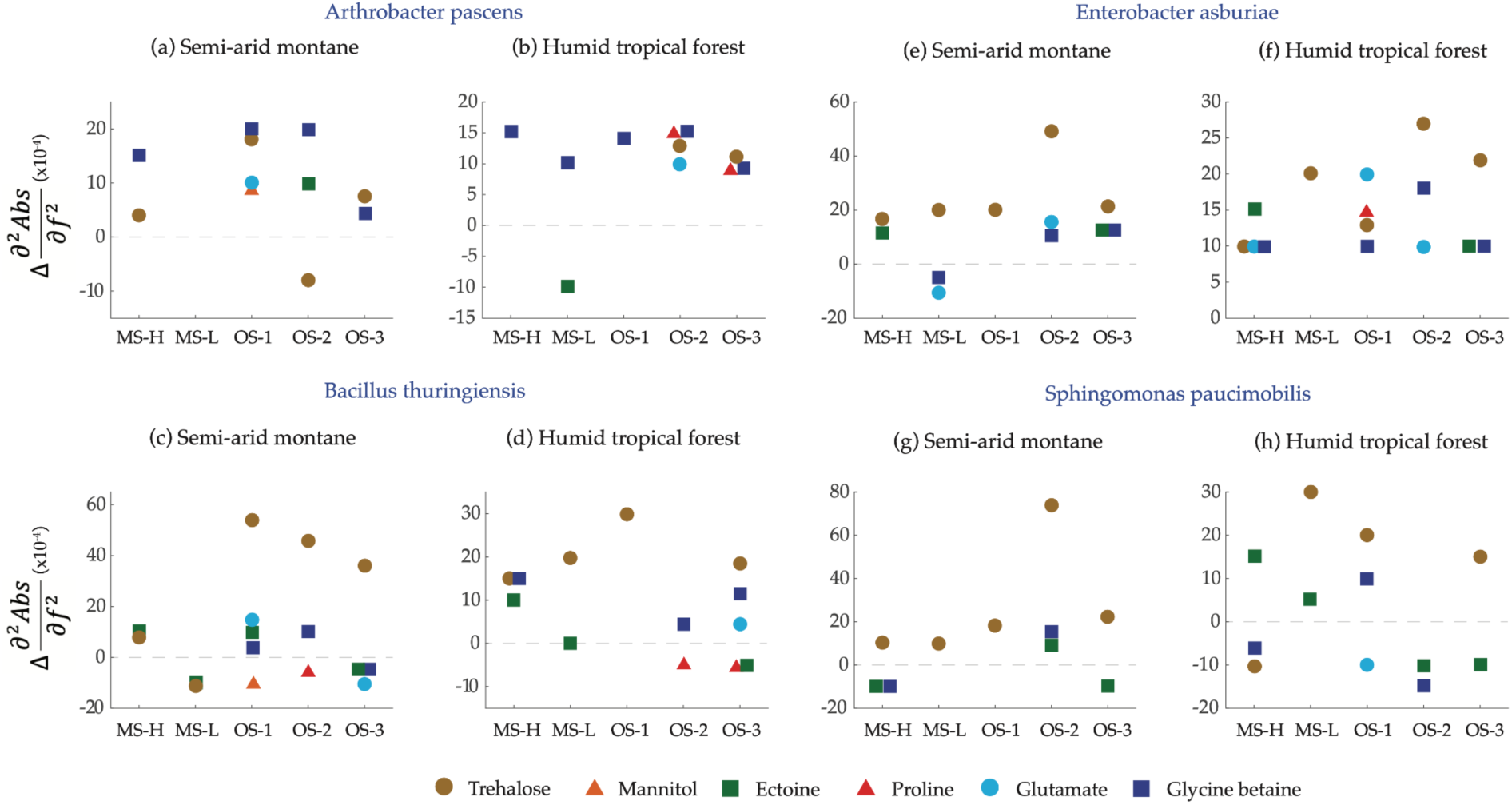
Changes in the relative concentration of different bacterial stress metabolites and osmolytes under matric and osmotic stress for each isolate. The general stress metabolites include trehalose, proline, and glutamate, while those associated with osmotic stress include mannitol, ectoine, and glycine betaine. These compounds have previously been shown to increase under drought conditions.

Doubling the NaCl concentration to 80 mM (OS3) provoked a strong elevation in the carbohydrate region for the HTF isolates (Fig. 6), a decline in the broader lipid region, and a generally negligible response in the υPO_2_^-^ region for most isolates. When considering distinct spectral features, the majority of both SAM and HTF isolates respond to the OS3 treatment through the accumulation of PHA (and in some cases PHB and polyphosphate), and the production of osmolytes. Trehalose increased in every isolate compared to control conditions (Fig. 7).

## Discussion

Soil microorganisms inhabit dynamic environments where rapid fluctuations in soil moisture are common^2^. During periods of hypo- and hyperosmolarity, prokaryotes must maintain positive intracellular turgor pressure to sustain growth and cell division^33^. The SR-FTIR spectromicroscopy approach determines the aggregate physiological responses and points to the metabolic trade-offs critical in maintaining diversity in microbial communities^34^. Our findings confirm the key role of osmolyte production under osmotic stress, while highlighting the importance of storage polymers and cell-cell signaling (υPO_2_^-^ region) as responses to stress. We synthesize this data below to address the overarching hypotheses of this study.

### Onset of matric and osmotic stress provokes contrasting metabolic responses in bacteria

Our initial hypothesis proposed that the different constituents of drought - matric stress and osmotic stress - will impart distinct metabolic responses in bacteria. Support for this hypothesis comes predominantly from the SAM isolates, which show distinct biochemical phenotypes between the matric and osmotic stress experiments (Fig. 4). An extensive analysis of the biochemical phenotype shows stress metabolites were more prominent under the OS experiments, while little response to the MS experiments was observed: Arthrobacter-SAM and Bacillus-SAM showed no clear deviation from the control phenotype under MS-L, while Sphingomonas and Enterobacter SAM isolates exhibited only moderate responses compared to the response to the OS treatments. These observations emphasize that organisms isolated from semi-arid soils are well acclimated to acute osmotic stress, reflecting their evolutionary adaptations to seasonal drought. In contrast, HTF isolates exhibited more similarity between the MS and OS experiments, while their stress responses rarely overlapping with the control conditions. This suggests a more restricted range of stress phenotypes within these isolates.

Bacteria adapt to fluctuations in their local environment through intracellular reallocation of resources to maintain cell integrity^39^, resulting in trade-offs between different pathways^40^. Our analysis highlights characteristic responses to stress observed in all isolates. The upregulation of cell signaling (represented by the υPO_2_^-^ region) was observed in most isolates under matric and osmotic stress experiments. These pathways commonly regulate bacterial response in dynamic environments^35^, preceding the general stress response of bacteria^36^, including the upregulation of the secondary metabolome^37^ and the production of biofilms^38^, all of which are important mechanisms mitigating the impact of osmotic stress on bacteria^2^.

Furthermore, conspicuous within the SR-FTIR data was the production and consumption of lipid storage polymers (i.e., PHAs, and PHBs) and glycogen, a carbohydrate storage compound, under matric and osmotic stress. These compounds are widely distributed amongst bacteria^41^, including the taxa from the present study^42–44^. The production and accumulation of storage polymers enable cells to survive prolonged periods of stress^45^. Starvation is a primary trigger upregulating the production of storage compounds^41^, facilitating the maintenance of metabolic activity when carbon availability declines^46^. Changes in osmotic potential under drought are often accompanied by a drop in substrate diffusion through soil pores^1^, resulting in cell starvation^2^. However, because our experiments were conducted under carbon replete conditions, the accumulation of storage compounds may result from a conditioned response to cells experiencing fluctuating osmotic potential. Alternatively, storage polymers are themselves important metabolic responses for protecting cellular integrity under osmotic stress^47,48^. Sedlack et al.^48^ noted that the production of PHA in *Cupriavidus* isolates decreased cytoplasmic membrane damage associated with osmotic shock. Similarly, PHB can enhance the survival of bacteria under both osmotic and oxidative stress conditions^47^.

In addition to the accumulation of energy storage polymers, our broad data analysis demonstrated evidence for the catabolism of lipid storage compounds concomitant with the production of carbohydrates (Supplementary Fig. 5). Lipid metabolism provides free fatty acids that can be used as a source of energy^49^, for carbohydrate production to maintain cellular integrity. However, it should be noted that such dynamics are potentially independent of one another: modeled output has demonstrated that the concurrent allocation to storage compounds and maintenance metabolism can be an optimal strategy under fluctuating environmental conditions^50^.

The third major response observed in the current experiments involves the production of polysaccharides, amino acids and osmolytes. Such upregulation is a common response within microbial isolates exposed to osmotic stress^16,17^, and soils undergoing drought^51,52^. We show here that osmolyte production is a common stress response for both semi-arid and tropical forest isolates (Fig. 7). The diversity of metabolites produced under the OS treatments tended to be higher than under the MS treatments, with multiple metabolites (e.g., trehalose, ecotoine, proline, and glycine betaine) produced under each of the OS treatments (Fig. 7), suggesting that more complex pathways of stress response are necessary under more stressful conditions.

The prominent role trehalose plays under the MS and OS conditions might be attributed to it being a general stress metabolite (Fig. 7). Trehalose production is commonly upregulated under osmotic or oxidative stress^5,53^ and heat shock^54^ and protects protein structure and membrane integrity. However, trehalose plays a multifaceted role in stress, also functioning as a storage compound^41^, and serves as an important substrate post-stress^55^. Given that compatible solute production is an energetically expensive process^56^, requiring the re-allocation or novel acquisition of resources to support synthesis, the upregulation of a versatile compound serves several requirements in a cells response to stress. Glycine betaine and ecotoine also played an important role in the overall stress response. In this case, both likely function as osmoprotectants, and are upregulated in response to changing osmotic potential^57^. For both Arthrobacter isolates glycine betaine was the primary metabolic response to stress, which suggests a stress response pathway conserved on evolutionary timescales, consistent with previous work on both bacteria and archaea^58^.

### Climate history will shape the metabolic response to stress

Our second hypothesis addressed the role climate history plays in determining the microbial response to stress. Climate history plays a critical role in shaping the adaptive capacity of microorganisms^9,22,59^, and we hypothesized here that isolates from semi-arid soils adapted to long seasonal drought would respond through a stronger metabolic response than isolates from humid tropical forest soils. The evidence supporting or refuting this hypothesis is complex. SAM isolates grow at a higher rate than HTF isolates under lower osmotic potentials (Fig. 2b), and maintained a biochemical phenotype similar to the control under matric stress (Fig. 4). They also showed a pronounced response to OS, with higher abundances of stress metabolites such as trehalose and ecotoine when compared to the HTF isolates (Fig. 7). Indeed, the climate-specific production of storage and stress compounds were consistently higher than control conditions for the SAM isolates, but not for the HTF isolates (Supplementary Fig. 5). Thus, this study highlights that SAM isolates, shaped by long-term drought legacies, show strong and more pronounced metabolic responses to stress that HTF, consistent with eco-evolutionary adaptation of microorganisms^60^.

We subsequently asked how pre-exposure to moderate stress would change the biochemical phenotype to more severe stress. Previous studies have shown bacteria to anticipate changes in their local environment, and respond accordingly^6,21^, and, in some cases, impart memory of a particular stimulus to descendants within a population^61,62^. In the current study, the OS2 experiment exposed cells to lower concentrations of NaCl prior to doubling the concentration. The overlap between OS2 and the other OS experiments (Fig. 4a,c,e,f,h) demonstrates a conserved response to osmotic stress, consistent with a general stress response^36^. However, in several cases the observed biochemical phenotype showed a response distinctive from that of either OS1 or OS3 (Fig. 4b,d,g). For Arthrobacter-HTF and Sphingomonas-SAM a stronger and more diverse metabolic response was noted for OS2 in both the stress metabolites (Fig. 7), and the storage compounds (Fig. 8), relative to OS1 and 3.

**Figure 8:**
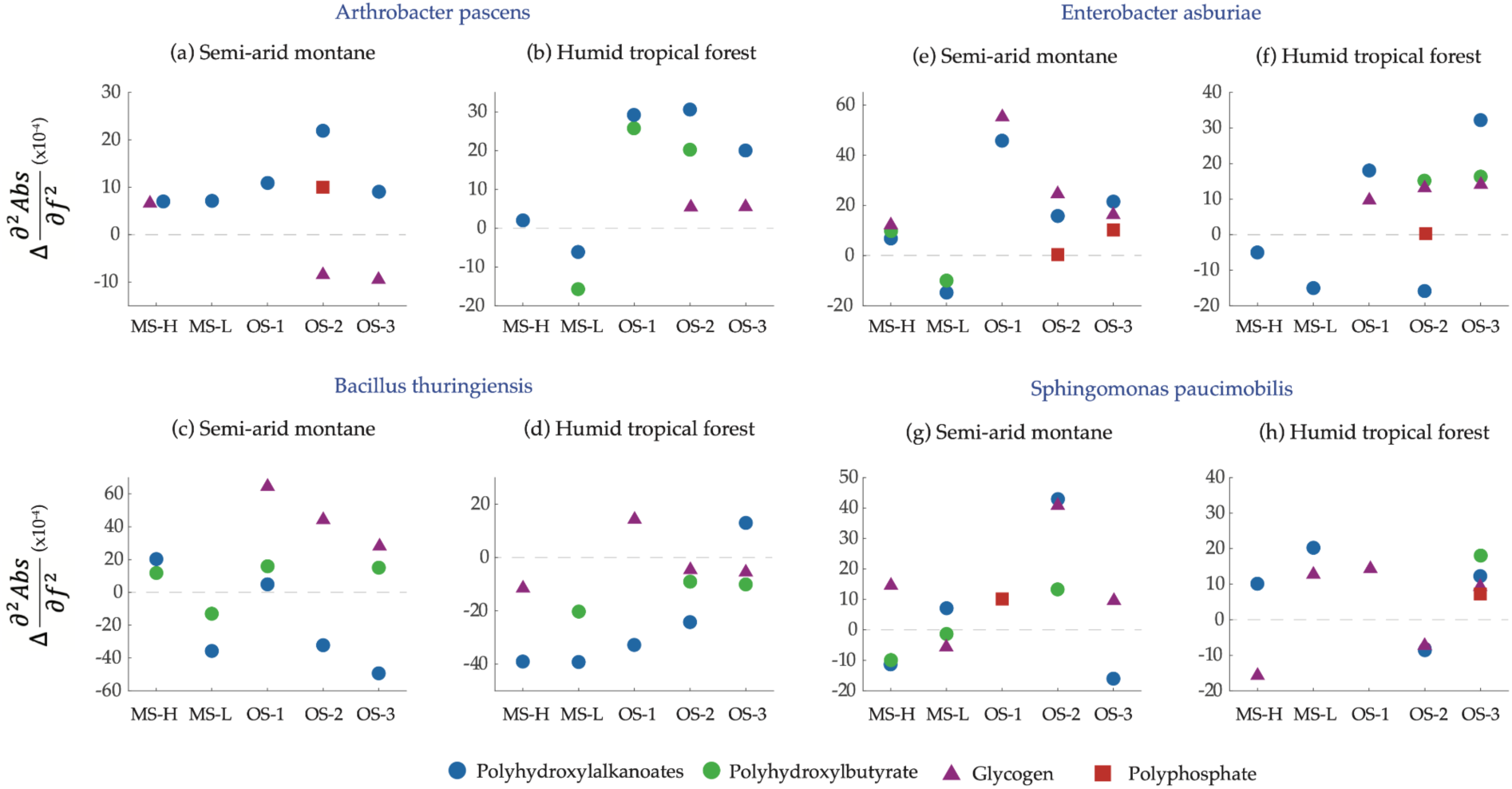
Specific metabolic shifts in energy storage molecules (polyhydroxyalkanoates, polyhydroxybutyrates, glycogen, and polyphosphate) under matric and osmotic stress.

## Conclusions

The current study characterizes the biochemical phenotype of bacteria isolated from semi-arid and tropical forest soils under matric and osmotic stress. We observed the response to stress is dependent on the nature of the stress, but also the climate history of the individual isolate, with SAM isolates showing greater adaptation to fluctuating environmental conditions. However, we recognize limitations with the current study. Most notably, the findings are limited to a small number of isolates that are represented at both sites. These findings are not, therefore, representative of the microbial community as a whole, which can show extremely divergent responses to soil moisture stress both within and between phyla^63^. Furthermore, cell-cell interactions, which increase under soil drying^12^, are likely as significant, if not more significant, in regulating the biochemical phenotype under stress. Maladapted populations could be outcompeted as environmental conditions change^64^ or deterministic interactions could lead to adaptation and co-existence between species^65^. Such interactions should be the focus of future research to determine how communities respond to a changing climate.

## Materials and Methods

### Isolations

Approximately 1 kg of soil was collected from a semi-arid montane (SAM) ecosystem within the East River Watershed, Colorado (38.92N, 106.9487W), and a humid tropical forest (HTF) on the Buena Vista Peninsula, Panama (9.185N, 79.8266W) within the Barro Colorado Nature Monument. The soil was sampled from the top 10 cm away from the plant roots. Select ecosystem properties are given in Table 1. Isolation of the soil bacteria proceeded according to previously published approaches^66^. Briefly, 10 g of soil was re-suspended and serially diluted ten-fold in a sterile saline solution (0.95 % NaCl). 0.1 ml of each dilution was plated onto one of six solid media, including various dilutions of R2A, R2G, DNBG, ISS, and the oligotrophic media, M9. The specific recipes for each media have been described previously^66^, and are provided in the supplementary material (Supplementary Table 2).

**Table 1:** Select edaphic parameters for the SAM site in Colorado and HTF site in Panama. *60 % of the mean annual precipitation falls as snow at this site. ^67^Hubbard et al., 2018, ^68^Sorensen et al., 2020, ^69^Bouskill et al., 2024, ^70^Paton et al., 2021, ^71^Chacon et al., 2023.

|  | East River Colorado<br>(Semi-Arid Montane) | Buena Vista Peninsula, Panama<br>(Humid Tropical Forest) |
| --- | --- | --- |
| **Climate** | Continental, subarctic<sup>67</sup> | Monsoonal<sup>70</sup> |
| **Soil Order** | Inceptisol<sup>68</sup> | Ultisol<sup>71</sup> |
| **Mean Annual Temp. (°C)** | 1<sup>67</sup> | 26<sup>70</sup> |
| **Mean Annual Ppt. (m)** | 1*<sup>67</sup> | 2.6<sup>70</sup> |
| **Relative Humidity (%)** | 45-67<sup>67</sup> | >80<sup>70</sup> |
| **Bulk Density** | 0.8 - 0.9<sup>68</sup> | 0.6 - 0.7<sup>71</sup> |
| **pH** | 6.7<sup>68</sup> | 5.7<sup>71</sup> |
| **Total Carbon (%)** | 6<sup>69</sup> | 5<sup>71</sup> |
| **Total Nitrogen (%)** | 0.2<sup>69</sup> | 0.43<sup>71</sup> |

The solid media was incubated for eight weeks, and colonies picked over time as they grew. To construct a diverse isolate library, we took an indiscriminate approach to colony picking, before serially diluting the colony in liquid media to ensure a pure culture. The lowest dilutions were re-plated on the initial media and regrown for 48 hours. Isolates were re-suspended in liquid media and grown overnight. A 0.5 ml aliquot of the axenic culture was vortexed with 0.5 ml glycerol and stored at -80 °C.

Glycerol stocks were all regrown within the same liquid media (M9: 0.16 M Na_2_HPO_4_, 0.11 M KH_2_PO_4_, 0.095 M NH_4_Cl, 0.043 M NaCl, 2x10^-3^ M MgSO_4_, 2x10^-4^ M CaCl_2_, 20 % glucose), to standardize media conditions for synchrotron experiments.

### Nucleic acid extraction and 16S rRNA sequencing

DNA was extracted from the axenic cultures using the *Quick*-DNA Fungal/Bacterial Miniprep kit (Zymo Research, CA, USA) according to the manufacturer’s protocol. 10 ng of genomic DNA was amplified in 25 μl PCR reactions containing 400 nM of each primer, 27F^72^ and 1492R^73^; and 2.5 U of DreamTaq Master Mix (Thermofisher Scientific, MA, USA). To obtain SSU rRNA gene sequences, PCR products were sequenced from both ends using primers 27F and 1492R at the UC Berkeley DNA Sequencing Facility (Berkeley, CA, USA). Templates were amplified using standard PCR conditions (95 ℃ x 5 mins, 30 cycles: 95℃, 52℃, 72℃ each for 30 seconds, 72℃ for 10 minutes).

A consensus sequence for each read was assembled (Geneious v10.11), and compared to the BLAST database to assign their taxonomic identity. The corresponding FASTA sequences were then aligned using a multiple sequence alignment program (MAFFT 7.0) and visualized in ITOL^74^. Further details of the isolates used in the present study are provided in Supplementary Table 1. For ease of reference, the isolates were named from both their broad genus name and the site of isolation (either SAM or HTF). For example, *Arthobacter Pascens* from the semi-arid montane site would become Arthobacter-SAM.

### Growth curves

The sensitivity of isolates to increasing NaCl concentrations (as a proxy for osmotic stress) was initially tested in batch culture. Isolates were grown in minimal media (M9 + 20 % glucose) with successively higher NaCl concentrations (2 - 750 mM), which can be converted to solute potential (Ψ_s_) using the following equation, Ψ_s_= -iCRT, where *i* is the molecules formed in water (2 for NaCl), C is the molar concentration of NaCl, R is pressure constant (= 0.0831 L·Pa·K^-1^·mole^-1^), and T is the ambient temperature (°K). Isolate growth (at OD 660 nm) was measured hourly over 72 hours and the specific growth rates were calculated during the exponential phase for each isolate under different NaCl concentrations using Monod kinetics^29^.

### Preparation of cells for SR-FTIR spectromicroscopy of metabolic responses to osmotic and matric stress

Cells were grown to the beginning of the exponential phase in M9 (with 20 % glucose), before being transferred to ^1^/_100_-strength M9 for attachment to silicon wafers. 4 μl of cells were placed onto a silicon wafer to promote attachment (Fig. 1a). The cells were washed three times in DI water to remove the unattached cells, and the remaining media. Ψ within this media at room temperature and atmospheric pressure was estimated to be ∼ 21 kPa. Subsequent experiments were designed to manipulate this value (Fig. 1b/c), allowing the biochemical response of microbes to changes in Ψ to be evaluated by SR-FTIR.

### Matric potential

Soil drying results in the formation of water films on soil particles, reducing diffusion and increasing the matric potential within the film. We replicated this process by allowing a solution of either ^1^/_5_ (termed MS-H) or ^1^/_10_ (MS-L) strength M9 media to dry down to 50 % of the original volume of 4 μl (fig. 1b). We estimated that this would increase Ψ from -220 kPa to -820 kPa under MS-H, and -120 kPa to -550 kPa under MS-L. This value is still within the estimated range of microbial activity during soil drying^15^.

### Osmotic potential

As soils dry, salts accumulate and present a significant physiological challenge to microbial cells. Osmotic shock under rapid soil drying can significantly impair microbial cell function, or provoke a metabolic response. Herein, we examined whether bacteria can mount a metabolic response to an acute osmotic shock by rapidly increasing NaCl in the media through the addition of either 40 mM NaCl (termed OS1, estimated Ψ ∼ -2.4 MPa), or 80 mM NaCl (OS3, estimated Ψ ∼ -3.6 MPa). We further asked whether pre-exposure to a lower concentration of NaCl can condition the cell upon subsequent acute exposure to a higher dose (80 mM). Previous work has shown that an immediate switch between a low and high dose of NaCl can invoke an anticipatory response under the higher dose^21^. This experiment, termed OS2, exposed the attached isolates to a 40 mM NaCl concentration for 20 minutes, before immediately increasing the NaCl concentration to 80 mM for an additional 20 minutes. For all experiments, the cells were washed in DI water and imaged under the FTIR microscope.

### SR-FTIR spectromicroscopy

All SR-FTIR spectromicroscopy measurements were carried out at the infrared beamline 1.4.3 of the Advanced Light Source (Lawrence Berkeley National Laboratory, Berkeley, CA; http://infrared.als.lbl.gov/) following previously published protocols^23,24^ Briefly, mid-infrared photons emitted from the synchrotron were focused on the monolayers of bacteria on the silicon-coated surface by the all-reflective optics infrared microscope (Figure 1d) equipped with a 32X, 0.65 numerical aperture objective. Each IR spectrum was raster scanned in the transflection mode using the IR microscope to map the bacterial monolayers at a 5 μm step size using Omnic 7.2 spectra software (Thermo Fisher Scientific, Waltham, MA). Each scan was repeated 64 times over the mid-infrared wavenumber range of 800–1800 cm^-1^, and each scan was recorded at a spectral resolution of 4 cm^-1^ with an adsorption peak position accuracy of ^1^/_100_ cm^-1^. Each SR-FTIR spectrum covered around 20 bacterial cells. All spectra for each sample map were exported for data pre-processing and multivariate analysis.

### SR-FTIR data pre-processing and chemometrics

The SR-FTIR spectra were subjected to data preprocessing. In-house scripts (Matlab, Mathworks, MA) were used to remove spectra from non-mono-layer distributed microbial cells or spectra with a poor signal-to-noise. Spectra that exhibited scattering effects were subject to the Kohler EMS correction algorithm to correct the spectral line shape distortion. The pre-processed spectra were then vector analyzed and transformed to the second derivative (7-point Savitzky-Golay smoothing, polynomial order 3) spectra. The 800–1800 cm^-1^ region was analyzed after mean-centering using principal component analysis (PCA) followed by linear discriminant analysis (LDA), which was performed on the outputs from PCA. Together, PC-LDA maximizes intergroup variance while minimizing the intragroup variance. The score plots of PC-LDA were used for a visual representation of the separation of the stress response in microbes from different geographic origins on the basis of their whole metabolic response to stress. To identify how different metabolic responses contribute to discrimination between samples we compared the mean adsorption spectra from the cluster loading plot. We focus our analysis here on the 900-1800 cm^-1^ fingerprint region that provides information on the production, transformation, or depletion of a range of compound classes, including the fatty acid carbonyl region (1680 – 1770 cm^-1^), protein secondary structure amide I (1600 - 1700 cm^-1^), amides II (1519 – 1544 cm^-1^) regions, and carbohydrate (1000-1300 cm^-1^) regions. Spectral interpretations were guided by the secondary derivative spectra and in-house reference spectra for polysaccharides, proteins, and water vapor. Quantitative identification of specific compounds related to bacterial storage compounds - polyhydroxyalkanoates (PHA), polyhydroxybutyrates (PHB), glycogen, polyphosphate - and metabolic stress compounds - trehalose, ectoine, glycine betaine, proline, glycogen - were identified through comparison of the secondary derivative spectra to the spectra for each compound (Supplementary Fig. 2b). The experimental spectra were analyzed for multiple characteristic peaks representing each compound (Supplementary Table 3), and an identification threshold set at 75 %.

### Flow cytometry

Live/dead staining was used to ascertain mortality rates under the OS and MS experiments. The OS and MS experiments described above were replicated in slightly larger volume (10 μl), and live/dead staining carried out following the manufacturers protocol (*BacLight* bacterial viability and counting kit, L34856, Molecular Probes, OR). Briefly, 10 μl control or treatment cells were combined with a 3.34 mM solution of SYTO 9 nucleic acid stain, and a 30 mM solution of propidium iodide, and incubated for 15 minutes at room temperature. This solution was passed through a cell sorter (Attune NxT Flow Cytometer, Thermo Fisher, Waltham, MA) to enumerate the proportion of live/dead cells post treatment.

### Data availability

In accordance with US-DOE data policy, the data presented in this manuscript will be deposited in the ESS-DIVE repository (https://ess-dive/gov) prior to publication.

## Acknowledgments

Funding for this work was provided by the US Department of Energy, Office of Science (BER), Early Career Research Program to N.J. Bouskill (#FP00005182). This research used resources of the Advanced Light Source, a U.S. DOE Office of Science User Facility under contract no. DE-AC02-05CH11231. We are grateful to Hans Betchel, who provided user support a ALS beamline 1.2 throughout this project.

