## Supplemental material for "Climate History and Metabolic Trade-offs shape the Bacterial Response to Drought"

### Supplementary Results

Under the high matric stress treatment (MS-H) the score plots showed a cluster (Fig. 3b) which exhibits a large degree of overlap among three tropical isolates the Gram-positive Bacillus-HTF, Arthrobacter-HTF, and the Gram-negative Sphingomonas-HTF. The accompanying vector plots (Supplementary Fig. 3c) show small differences among these three tropical isolates, lower absorbance in the 1770 - 1720 $cm^{-1}$ spectral region of lipids, 1570-1510 $cm^{-1}$ of amide II, 1490 - $CH_2$ and $CH_3$ and higher absorbance in the $\upsilon PO_2^{-}$ (~1250-1200 $cm^{-1}$) region. These overlapping tropical isolates are complimented by three adjacent semi-arid isolates, the Gram-negative Enterobacter-SAM, Sphingomonas-SAM, and the Gram-positive Arthrobacter-SAM. The cluster vectors of Enterobacter-SAM and Arthrobacter-SAM show lower absorbance at ~1730 $cm^{-1}$ in the lipid region and much higher absorbance at ~1110 $cm^{-1}$ and ~1076 $cm^{-1}$ in the carbohydrate region. In contrast, the cluster vector of Sphingomonas-SAM shows lower absorbance at ~1730 $cm^{-1}$ but much higher absorbance at 1214 $cm^{-1}$, similar to Sphingomonas-HTF. The final two isolates, Bacillus-SAM, which split into two subgroups, and Enterobacter-HTF, ordinate away from the main cluster and each other. The most dominant wavenumbers in distinguishing Bacillus-SAM and Enterobacter-HTF from the other isolates are the large absorbance at ~1730 $cm^{-1}$ and ~1155 $cm^{-1}$ respectively. Moreover, the vector spectra show that the discrepancy between the two Bacillus isolates from different climate ecosystems lies in divergent responses in the lipid/fatty acid, proteins, and carbohydrate regions, with prominent increases at 1730, 1664, 1490-1390, 1295, 1135, 1050-1010 $cm^{-1}$ for Bacillus-SAM, and decreases at 1710, 1598, 1215, and 1076 $cm^{-1}$. Most of the difference between Enterobacter-HTF and Enterobacter-SAM can be explained by differences in the carbohydrate (1250-1100 $cm^{-1}$) region, with important contributions from amide I /II.

Even under the milder MS experiment (MS-L), we observed a similar cluster of tropical isolates (Arthrobacter-HTF, Bacillus-HTF, and Enterobacter-HTF), alongside the phylogenetically similar semi-arid counterparts for Arthobacter-SAM and Enterobacter-SAM (Fig. 3c). The cluster vector plots (Supplementary 3c) show small differences among these tropical isolates, or the semi-arid isolates except for the higher absorbance at ~1648 $cm^{-1}$, ~1496 $cm^{-1}$, ~1400 $cm^{-1}$, and ~1223 $cm^{-1}$ in the Arthrobacter-SAM isolate. The Sphingomonas isolates ordinate away from this large cluster, and directly opposite one another. This implies unique spectral responses to treatment between these two isolates, with significant contributions from Sphingomonas-HTF and Sphingomonas-SAM at ~ 1102 $cm^{-1}$ and ~1020 $cm^{-1}$ (carboxylates) respectively. The Bacillus-SAM ordinated away from the Bacillus-HTF (and all isolates), predominantly due to major contributions from the high lipid peak at ~1740 $cm^{-1}$ and the double carbohydrate peaks at ~1136 $cm^{-1}$ and ~1028 $cm^{-1}$, and the low absorbance at ~1630 $cm^{-1}$ (protein amide I), ~1225 $cm^{-1}$, and ~1075 $cm^{-1}$.

The clustering emerging under osmotic stress experiments was related to both phylogeny and climate history (Fig. 3d, e, f). Climate history tended to guide the biochemical response under direct exposure (i.e., OS1 & OS3), while the step-up experiment (OS2) with priming "osmotic stress" resulted in a response clustering by phylogeny. A strong cluster of multiple isolates emerged under the milder OS experiment (OS1). Furthermore, the most severe OS experiment (OS3) also showed evidence for the emergence of unique biochemical phenotypes for each isolate, with the HTF cluster more diffuse under this condition (Fig. 3f).

We found that bacterial exposure to osmotic stress resulted in fundamentally different changes (relative to matric stress and the stress-free controls) in cell biochemistry. Under OS1, the score plots (Fig 3d) and cluster vector spectra (Supplementary Fig. 3a) illustrate a large degree of co-clustering and overlap. The tropical isolates Arthobacter-HTF, Bacillus-HTF, and Sphingomonas-HTF showed biochemical response similarity (Fig. 3d), with showing higher amplitude peaks at ~1724 $cm^{-1}$, ~1710 $cm^{-1}$, ~1593 $cm^{-1}$, ~1216 $cm^{-1}$, ~1186 $cm^{-1}$, ~1063 $cm^{-1}$, and ~1056 $cm^{-1}$ in Arthrobacter-HTF. The semi-arid isolates, Bacillus-SAM and Enterobacter-SAM showed overlapping peaks with little differences. Similarly, Arthrobacter-SAM and Sphingomonas-SAM co-clustered, however, the amplitude peaks of Arthrobacter-SAM at ~1730 $cm^{-1}$, ~1710 $cm^{-1}$, ~1690 $cm^{-1}$, ~1498 $cm^{-1}$, ~1225 $cm^{-1}$ and ~1125 $cm^{-1}$ were larger. Finally, the tropical Enterobacter-HTF ordinated alone along the primary axis which accounted for most (80%) of the variations, and the secondary axis which accounted for 9% of the variations. This distinct separation was defined by a significant absorption peak at ~1110 $cm^{-1}$ (Supplementary Fig. 3c).

A subsequent priming OS experiment (OS2), that pre-conditions the isolates to the lower osmotic stress (i.e., OS1) before exposure to higher stress, revealed the emergence of a co-cluster of bacterial isolates showing similar biochemical responses to the step-wise experiment (Fig. 3e). This large co-cluster again showed similarity in the responses of phylogenetically similar organisms isolated from different environments, containing both the tropical isolates Bacillus-HTF Sphingomonas-HTF and semi-arid isolates Bacillus-SAM, Enterobacter-SAM, and Arthrobacter-SAM, the latter was split into two small subgroups. The cluster vectors of Bacillus-HTF, Enterobacter-HTF, and Arthrobacter-SAM in Supplementary Fig. 3c showed overlapping peaks at ~1466 $cm^{-1}$, ~1444 $cm^{-1}$, ~1230 $cm^{-1}$, ~1221 $cm^{-1}$, and ~1120 $cm^{-1}$. Enterobacter-HTF and Arthrobacter-SAM showed overlapping peaks at ~1523 $cm^{-1}$ (amide II), ~1225 $cm^{-1}$ ($\upsilon PO_2^-$), ~1100 $cm^{-1}$, and ~1068$cm^{-1}$ (carbohydrates). The important feature in distinguishing Arthrobacter-HTF from this 6-member cluster along the secondary axis (PC-LDA2) was the large amplitudes of peaks at ~1466 $cm^{-1}$, ~1444 $cm^{-1}$, ~1230 $cm^{-1}$, ~1221 $cm^{-1}$, and ~1120 $cm^{-1}$. Sphingomonas-SAM differed from all others across the primary axis (PC-LDA1), which accounted for 69% of the variations. The vector plot showed a large difference between Sphingomonas-SAM and all other isolates within the lipid region (~1745 $cm^{-1}$), amide I (~1662 $cm^{-1}$, ~1630 $cm^{-1}$, ~1625 $cm^{-1}$), $\delta_{as}$ and $\delta_s$ (C–H) of $CH_3$ and $CH_2$ in lipids and proteins (~1488 $cm^{-1}$, ~1440 $cm^{-1}$, ~1364 $cm^{-1}$, ~1337 $cm^{-1}$, ~1326 $cm^{-1}$, ~1316 $cm^{-1}$, and carbohydrates (~1180 $cm^{-1}$, ~1163 $cm^{-1}$, ~1154 $cm^{-1}$, ~1066 $cm^{-1}$, ~1053 $cm^{-1}$).

Finally, relatively unique biochemical responses emerged under the most severe OS experiment (OS3). The Enterobacter isolates show some overlap in their response, but this might be attributed to a heterogeneous response of different microcolonies of Enterobacter-HTF, which shows clear variability rather than a succinct cluster. In general, the response to OS3 produces strong heterogeneity across the isolates as they respond to rapid changes in external osmotic pressure (Fig. 3f). Here the semi-arid Gram-negative Sphingomonas-SAM and the two tropical Gram-positive isolates Arthrobacter-HTF and Bacillus-HTF differed from the others along the primary PC-LDA axis that accounted for 65% of the variation in the OS3 data, which was largely defined by the absorbance in amide I and II (~1694 $cm^{-1}$, ~1653 $cm^{-1}$, ~1635 $cm^{-1}$, 1548 $cm^{-1}$), $\delta_{as}$ and $\delta_s$ (C–H) of $CH_3$ and $CH_2$ in lipids and proteins (~1427 $cm^{-1}$, ~1440 $cm^{-1}$, ~1400 $cm^{-1}$, ~1364 $cm^{-1}$), $\upsilon PO_2^-$ (~1248 $cm^{-1}$, ~1080 $cm^{-1}$), and carbohydrates (~1160 $cm^{-1}$ , ~1150 $cm^-$

$^{1}$, ~1100, ~1065, and 1024). Sphingomonas-SAM and -HTF differed from other isolates along the second PC-LDA axis, which accounted for 19% of the variation, and was defined by the absorbance in ~1510 $cm^{-1}$, ~1220 $cm^{-1}$, ~1155 $cm^{-1}$, ~1105 $cm^{-1}$, and ~1050 $cm^{-1}$. Overall, the SAM isolates orientated around a larger cluster of tropical isolates, with strong separation between the SAM isolates. By contrast, the clustering of HTF isolates would suggest a similar response to OS3, however, the wider cluster of replicate cells for each isolate would further indicate within-population heterogeneity in response.

**Supplemental material**

Supplemental figures for the main text.……………………………………………3

Supplemental tables for the main text…………………………………………...18

### Supplemental Figures

**Supplemental Figure 1**: Survival rates of all isolates under the MS and OS treatments. Survival is based on live-dead staining and flow cytometry.

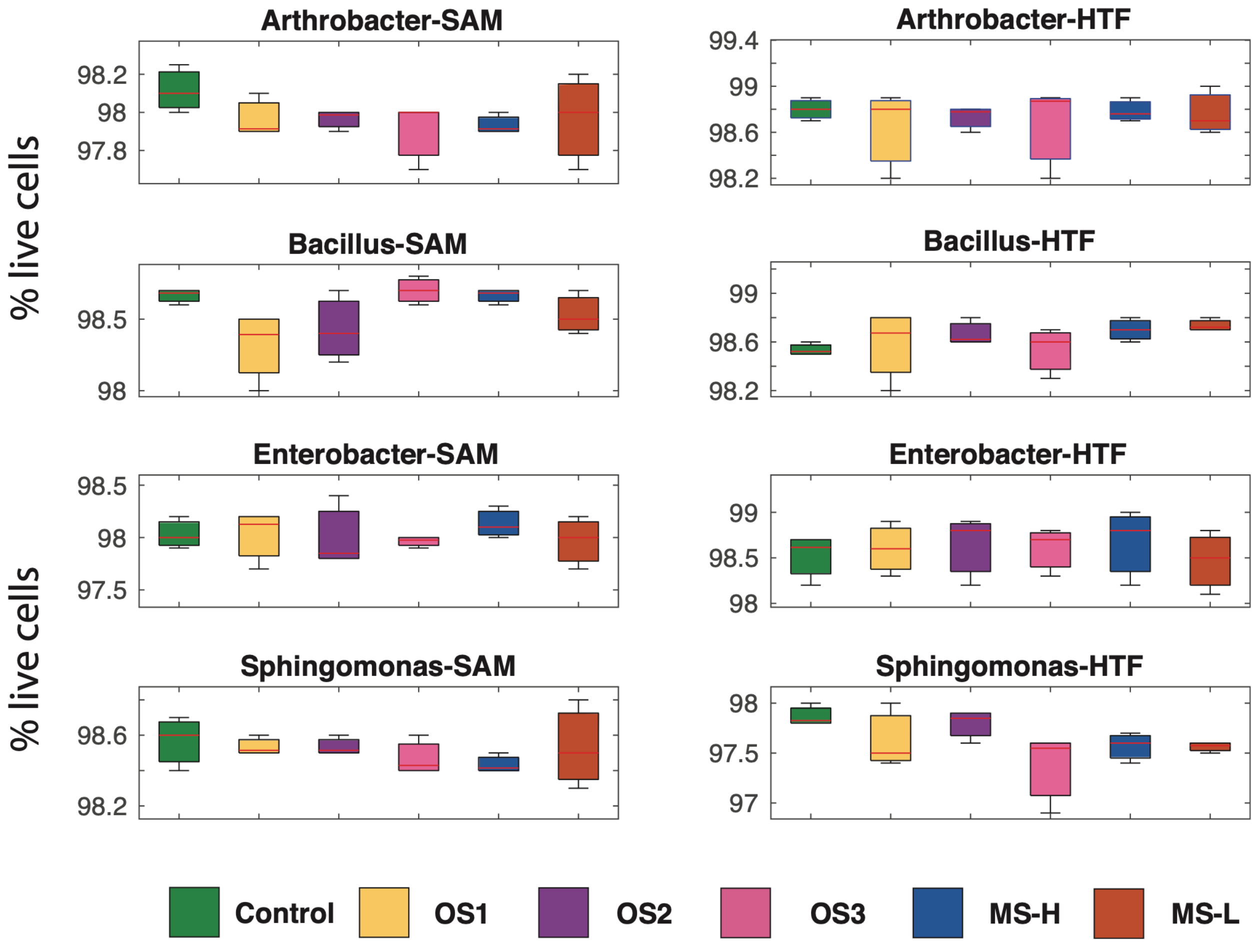

**Supplemental Figure 2a:** Characteristic compound absorbance spectra for fatty acids, proteins, and carbohydrates. The bottom spectrum (-) is an example of a mean cluster absorbance spectrum (see main text), while the top spectrum (-) shows its secondary derivative spectrum which is used to resolve overlapping peaks in the data set.

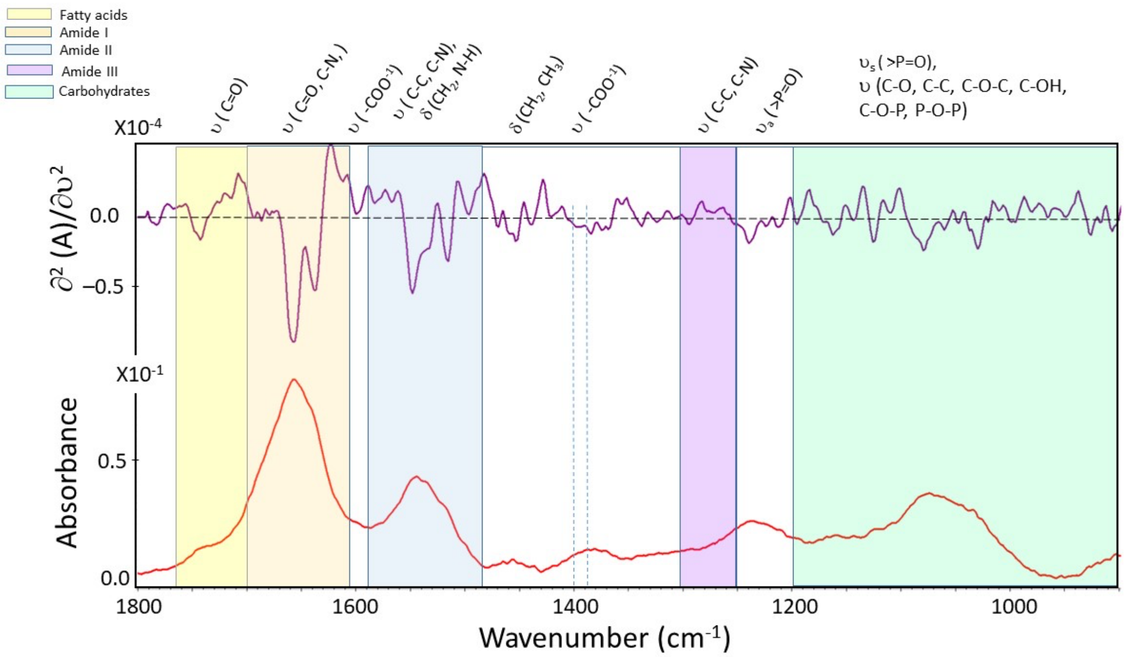

**Supplemental Figure 2b:** Characteristic compound absorbance spectra for osmolytes involved in microbial stress response. Characteristic peaks for each osmolyte are provided in Supplemental Table 3.

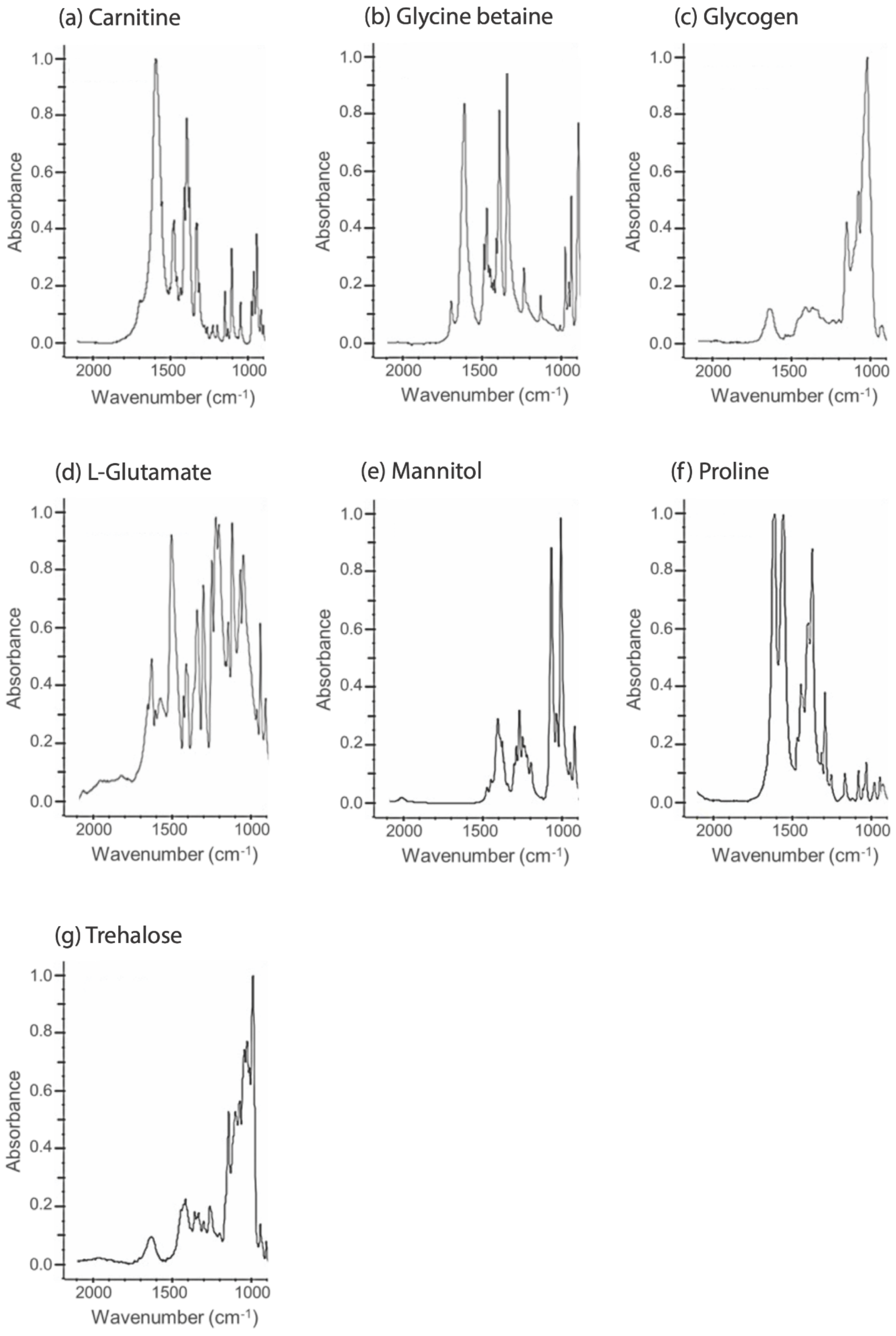

**Supplemental Figure 2c:** Individual 2nd derivatives of the mean spectra for each bacterial isolate. Each panel represents a different experimental condition as follows: (a-b) Matric stress-H/-L, (c,d,e) Osmotic stress 1,2,3., and (f) controls. The negative peaks of the 2nd derivative spectra are used to resolve overlapping peaks.

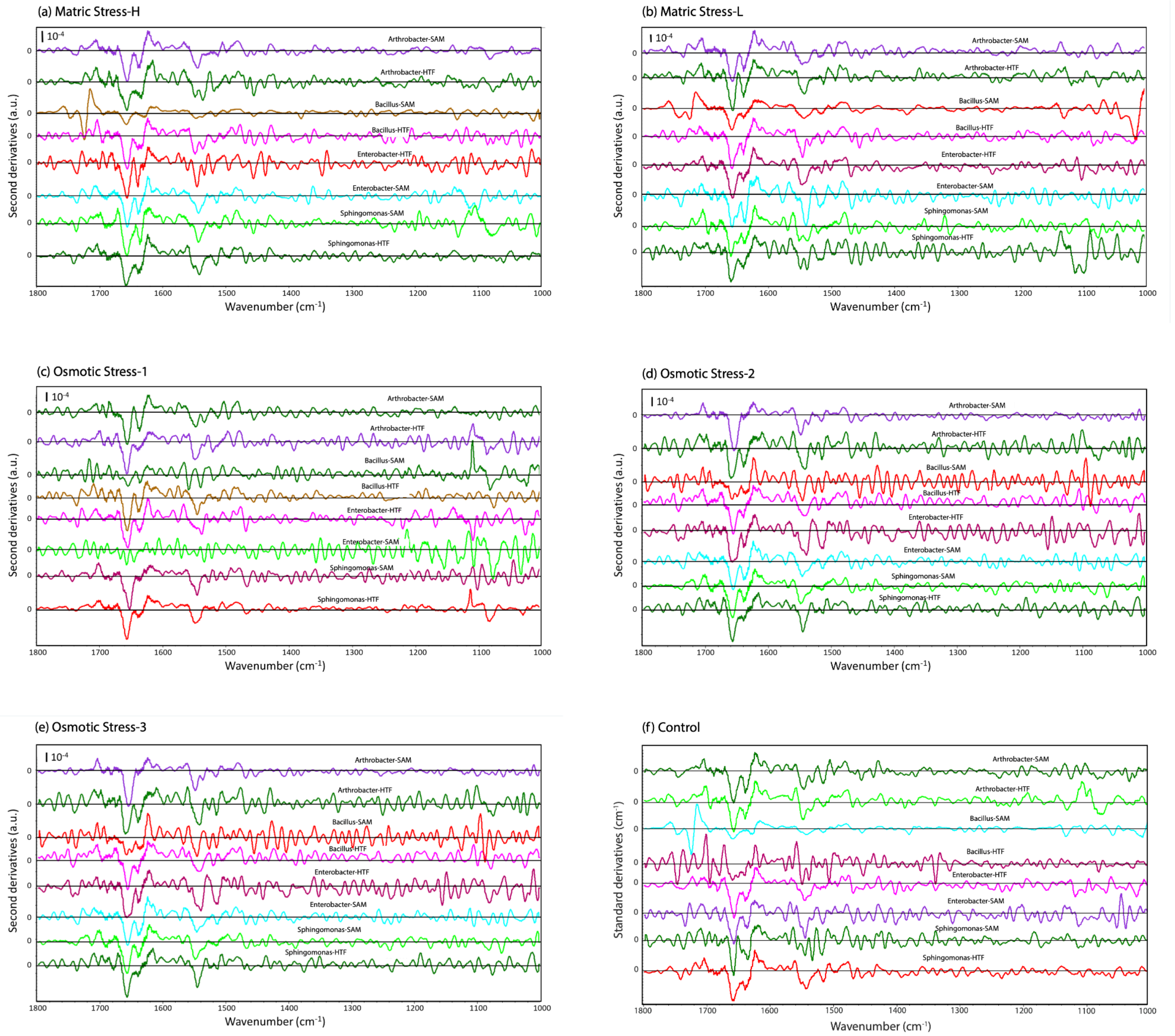

**Supplemental Figure 3a:** The cluster vector spectral plots depicted as the difference between each bacterial isolate as compared with the mean spectrum (plotted as the baseline) at different experimental conditions as follows: (a) control, (b,c) Matric stress-H/-L, (d-f) Osmotic stress 1,2,3. All cluster vector spectra are shifted for ease of visualization.

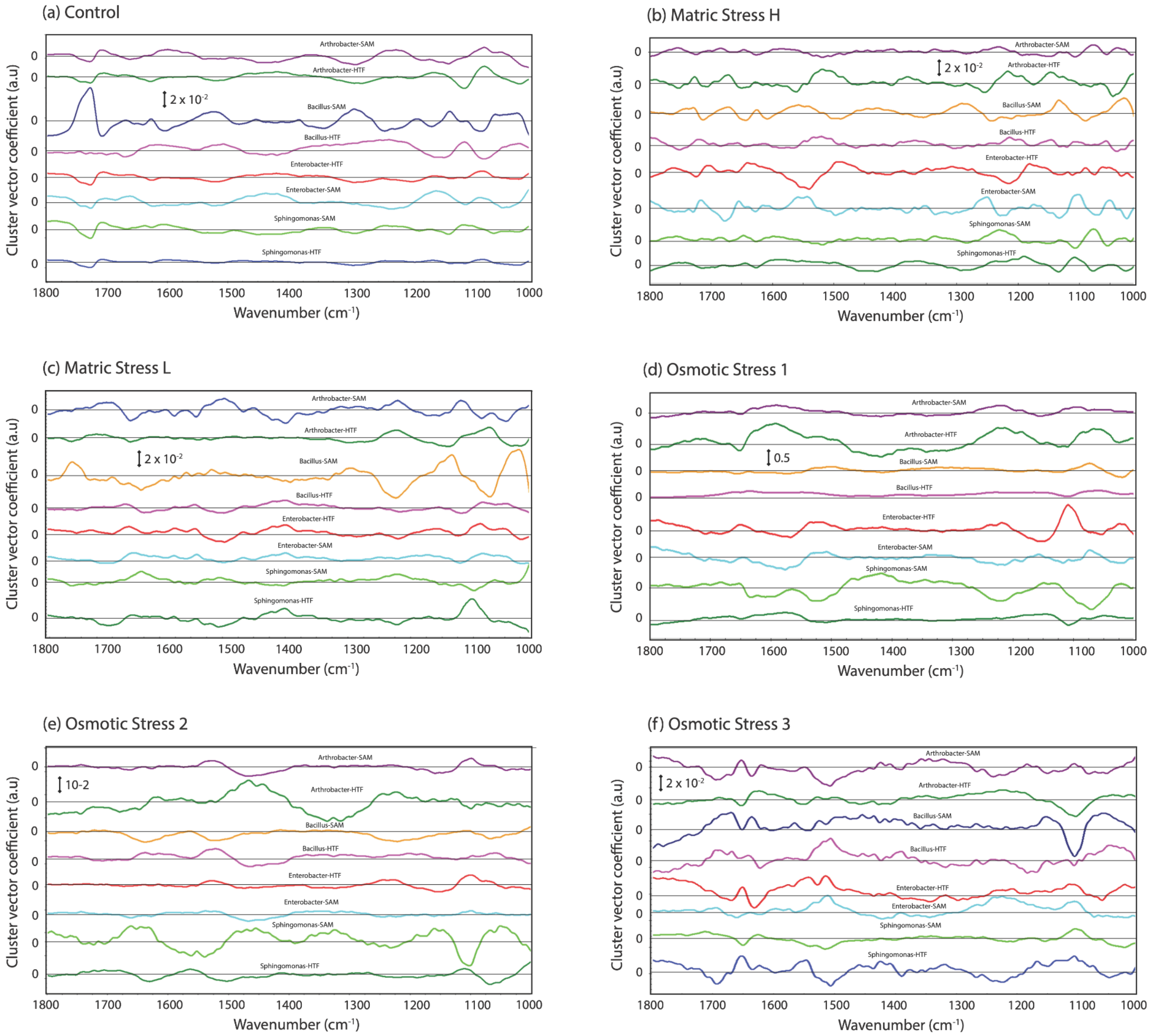

**Supplemental Figure 3b:** The first three PC-LDA loading plots. These plots are based on all relative positioning of all isolates together (i.e., Arthrobacter-SAM, Arthrobacter-HTF, Bacillus-SAM, Bacillus-HTF, Enterobacter-SAM, Enterobacter-HTF, Sphingomonas-AM, and Sphingomonas-HTF) under different experimental conditions as follows: (a) control, (b,c) Matric stress-H/-L, (d-f) Osmotic stress 1,2,3.

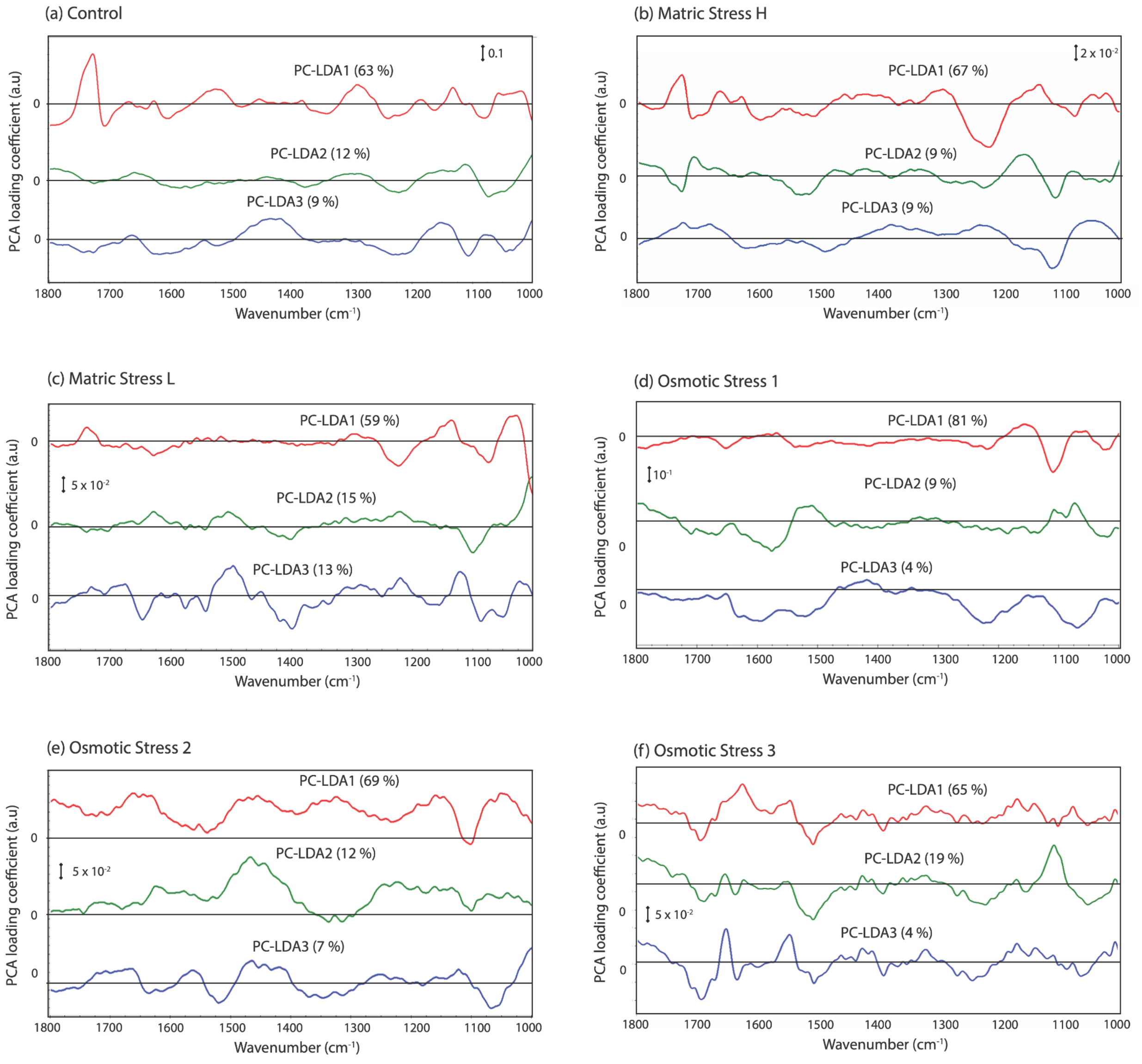

**Supplemental Figure 3c:** Cluster vector plots depicted as the difference between each bacterial isolate as compared with the mean spectrum (plotted as the baseline) at different experimental conditions as follows: (a,b) Matric stress-H/-L, (c-e) Osmotic stress 1,2,3. All cluster vector spectra are plotted with a common scale.

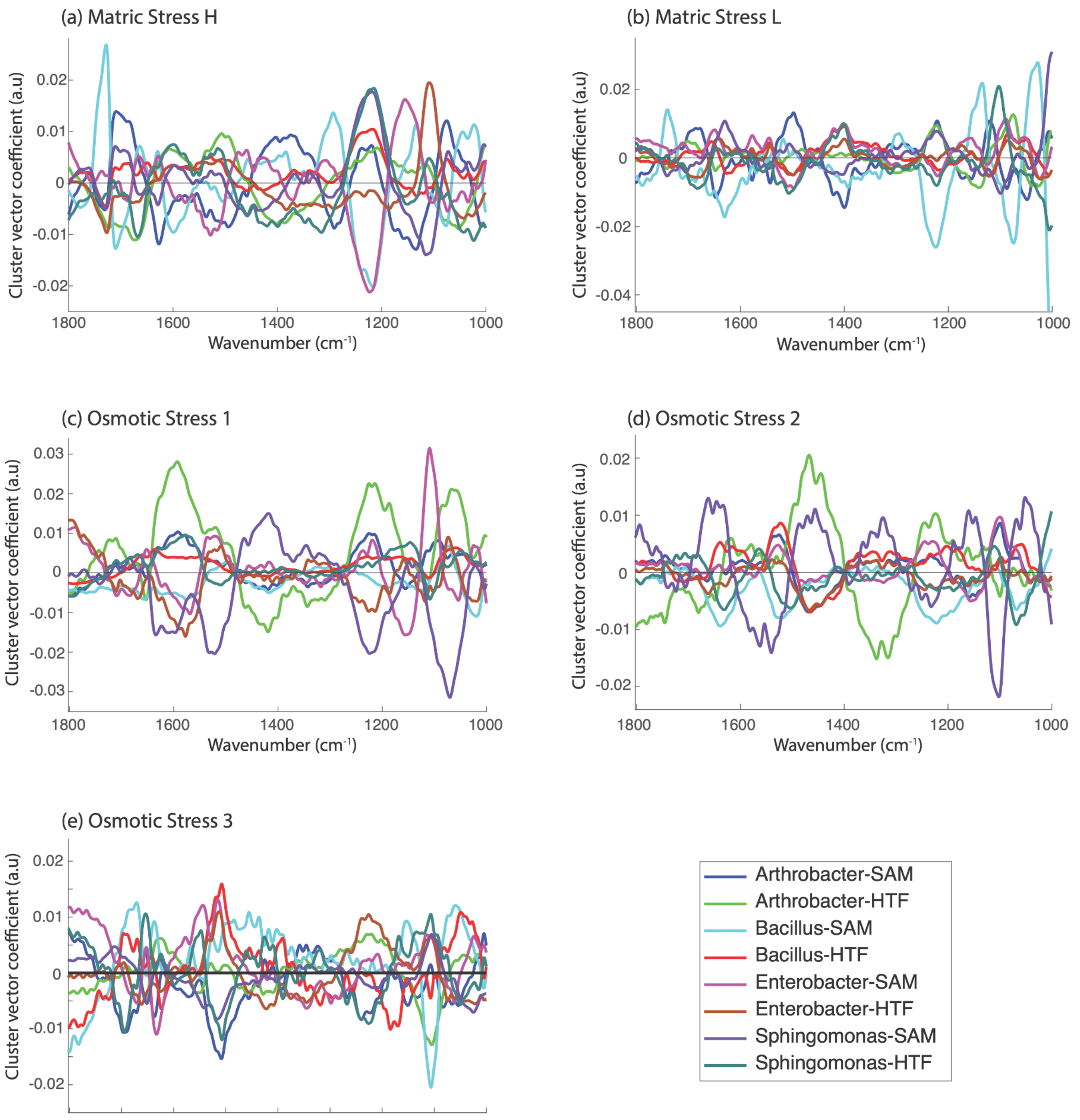

**Supplemental Figure 3d:** The cluster vector spectral plots depicted as the difference between each experimental condition as compared with the mean spectrum (plotted as the baseline) for different bacterial isolates as follows: (a) Arthrobacter-SAM, (b) Arthrobacter-HTF, c) Bacillus-SAM, (d) Bacillus-HTF, (e) Enterobacter-SAM, (f) Enterobacter-HTF, (g) Sphingomonas-AM, (h) versus Sphingomonas-HTF. All cluster vector spectra are plotted with a common scale.

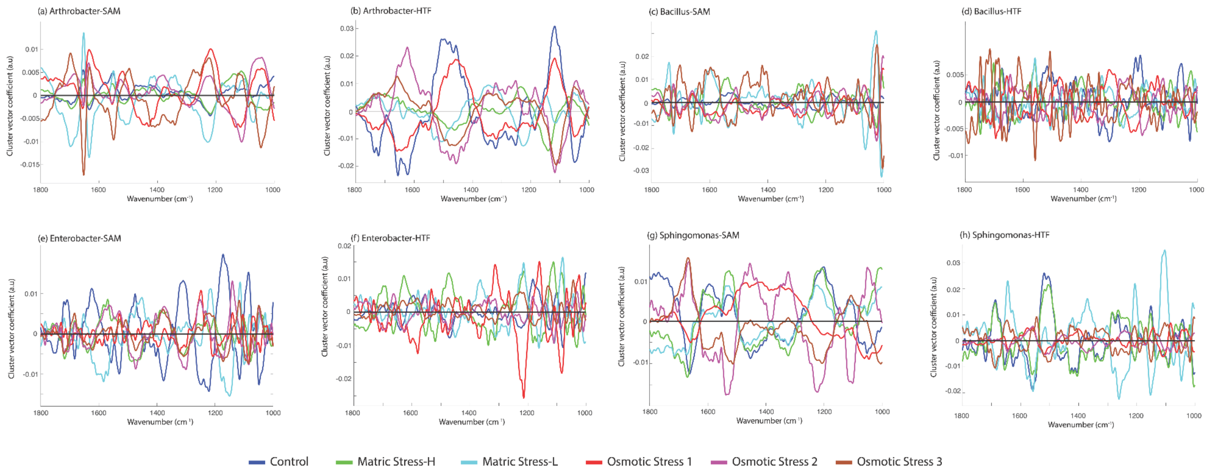

**Supplemental Figure 4a:** Ordination plots across the first three factors based on Principal Component-Linear Discrimination Analysis for each isolate. Each panel, visualized with Varimax Rotation, shows the relative dissimilarity of a single isolate under different experimental conditions of osmotic and matric stress. The panels provide results for bacteria isolated from semi-arid montane (SAM) and humid tropical forest (HTF).

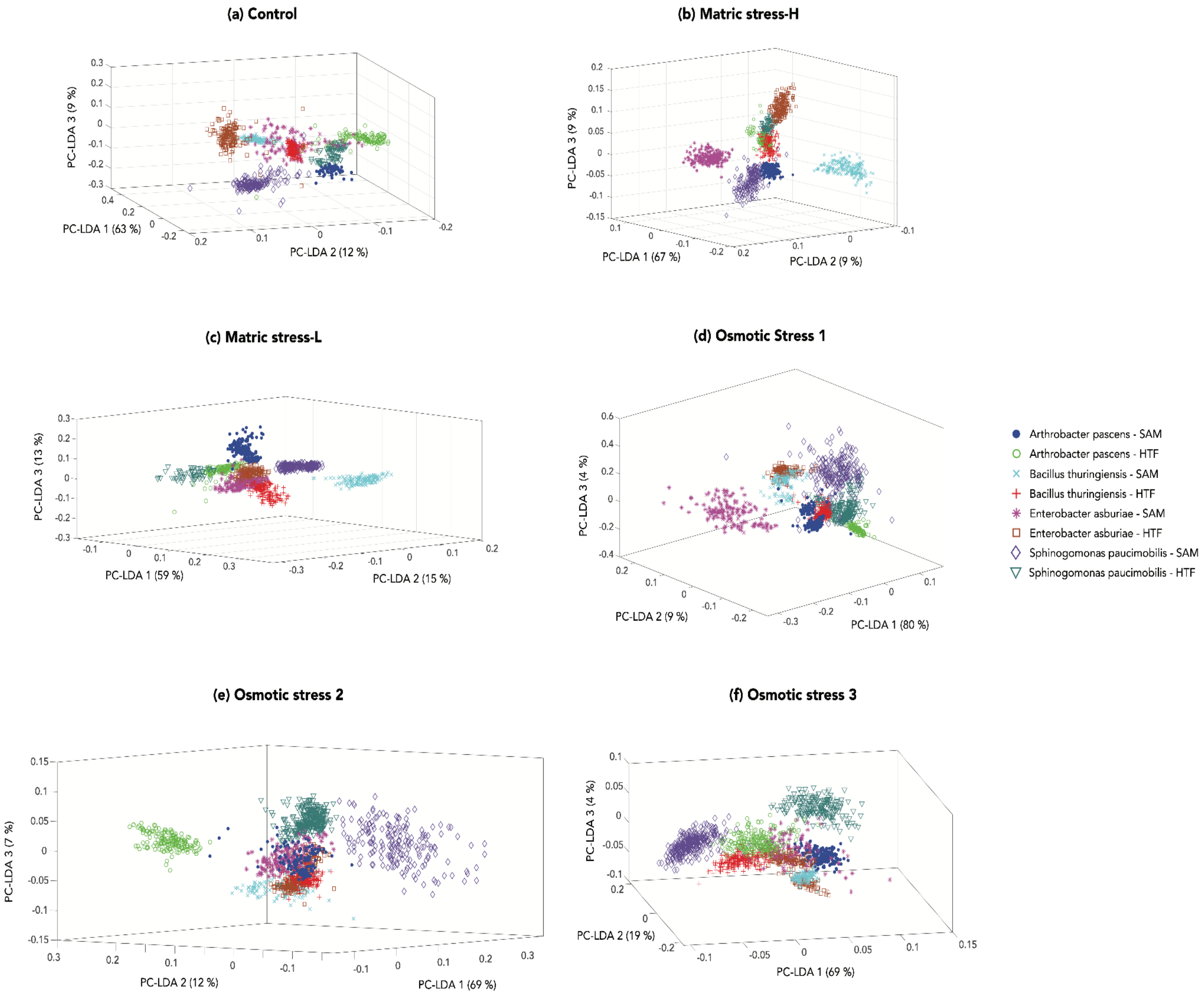

**Supplemental Figure 4b:** The first three PC-LDA loading plots of controls versus matric stress-H versus matric stress-L versus osmotic stress 1 versus osmotic stress 2 versus osmotic stress 3 for each isolate as follows: (a) Arthrobacter-SAM, (b) Arthrobacter-HTF, (c) Bacillus-SAM, (d) Bacillus-HTF, (e) Enterobacter-SAM, (f0 Enterobacter-HTF, (g)  Sphingomonas-AM, and (h) Sphingomonas-HTF.

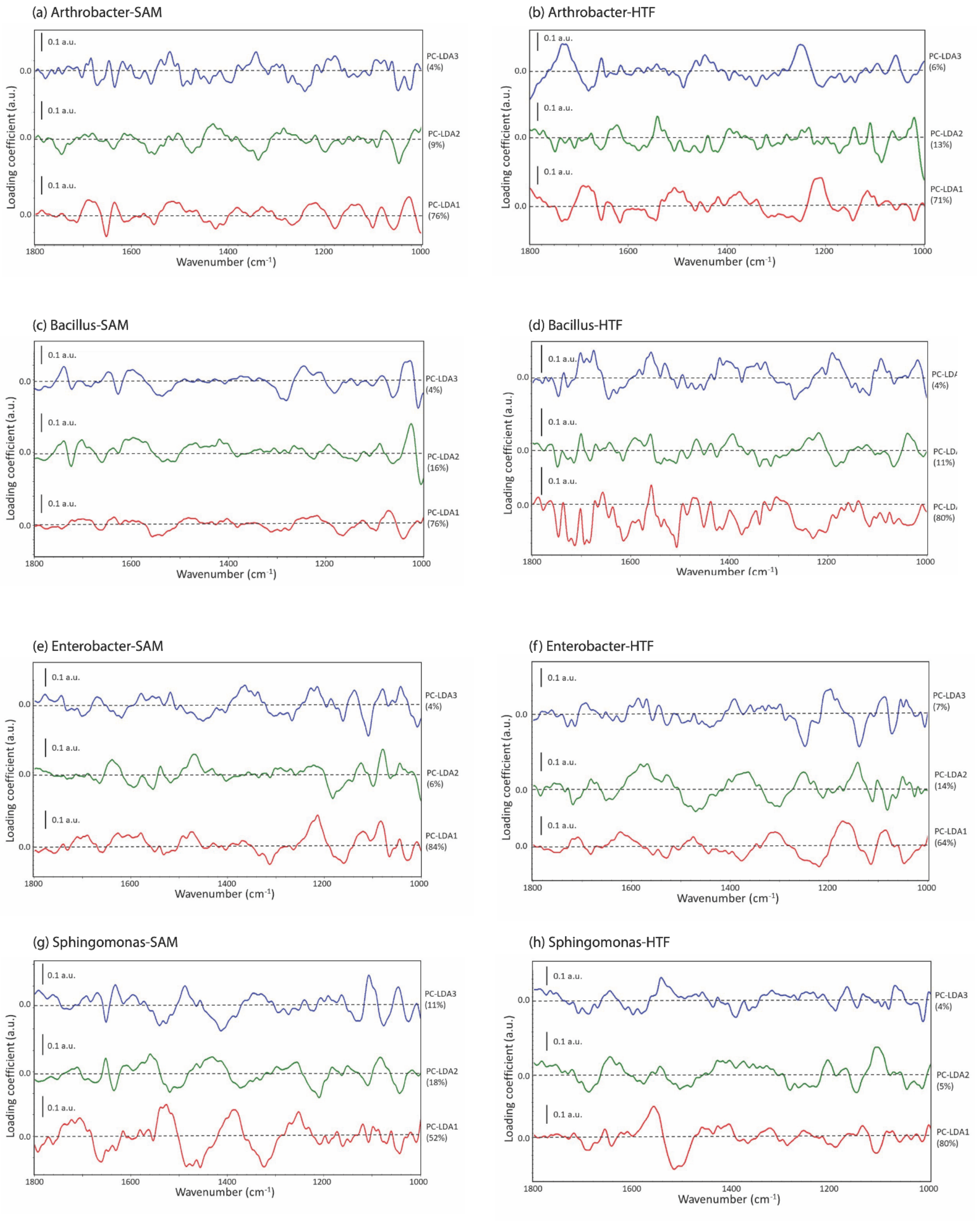

**Supplemental Figure 5:** Changes in the abundance of (a) storage compounds (e.g., PHA, PHB, glycogen), and (b) osmolytes (e.g., trehalose, glycine betaine, etc.) relative to the control conditions. The data has been aggregated by climate history and experiment (e.g., MS-H, OS-1, etc.). Each color represents a different experiment with SAM: opaque shading, and HTF: transparent shading.

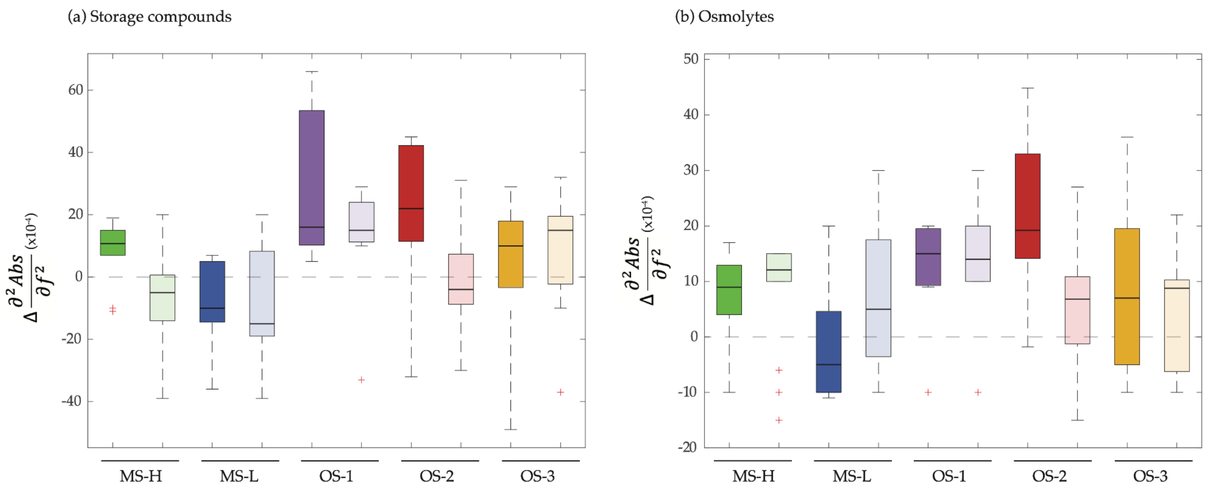

**Supplemental Tables**

**Supplemental Table 1**: Isolates used in the current study, location of isolation, and GenBank accession number.

| Isolate name | Study ID* | Isolation location | Accession number** |
|---|---|---|---|
| *Arthrobacter pascens* | Arthrobacter-SAM | Colorado | OQ566217 |
| *Arthrobacter pascens* | Arthrobacter-HTF | Panama | OQ566218 |
| *Bacillus thuringiensis* | Bacillus-SAM | Colorado | OQ566219 |
| *Bacillus thuringiensis* | Bacillus-HTF | Panama | OQ566220 |
| *Enterobacter asburiae* | Enterobacter-SAM | Colorado | OQ566221 |
| *Enterobacter asburiae* | Enterobacter-HTF | Panama | OQ566222 |
| *Sphingomonas paucimobilis* | Sphingomonas-SAM | Colorado | OQ566223 |
| *Sphingomonas paucimobilis* | Sphingomonas-HTF | Panama | OQ566224 |

*SAM: Semi-Arid Montane; HTF: Humid Tropical Forest.
**Sequences deposited in GENBANK

**Supplemental Table 2:** Isolation media used in the current study.

*R2A/ R2G*

| | R2A (1/10 strength) (g)* | R2G (1/20 strength) (g)* |
|---|---|---|
| Protease Peptone | 0.05 | 0.025 |
| Starch | 0.05 | 0.025 |
| Glucose | 0.05 | 0.025 |
| Yeast Extract | 0.05 | 0.025 |
| Casein hydrolysate | 0.05 | 0.025 |
| Dipotassium phosphate | 0.3 | 0.3 |
| Sodium pyruvate | 0.3 | 0.15 |
| Magnesium sulfate | 0.024 | 0.024 |
| Sodium chloride | - | 4.5 |
| Agar | 14 | - |
| Gelrite | - | 12 |

*DNGB*

| | DNGB (g)* |
|---|---|
| Difco nutrient broth | 0.08 |
| Calcium Chloride | 0.07 |
| Gellan gum | 8 |

*Inorganic salt starch media*

| | **ISS (g)*** |
|---|---|
| Starch | 10 |
| Dipotassium phosphate | 1 |
| Magnesium sulfate | 1 |
| Sodium chloride | 1 |
| Ammonium sulfate | 2 |
| Calcum carbonate | 2 |
| Ferrous sulfate | 0.001 |
| Manganese chloride | 0.001 |
| Zinc sulfate | 0.001 |
| Agar | 20 |

*1L with ultrapure water (pH = 7.2)

**Supplemental Table 3:** Characteristic absorption peaks for storage and osmotic stress metabolites.

| Compounds | IR absorption peaks ($cm^{-1}$) | References |
|---|---|---|
| **Energy Storage compounds** | | |
| Polyhydroxyalkanoates (PHA) | ~1745-1710 (~1744, ~1740, ~1732, ~1728, ~1720), ~1391, ~1282, ~1262, ~1257, ~1178, ~1128, ~1094, ~1015 | Christensen et al 2023; Ojha & Das 2020; Hu et al. 2013; Arcos-Hernandez et al, 2010; Misra et al. 2000; |
| Polyhydroxybutyrates (PHB) | ~1732, ~1379, ~1278, ~1183, ~1132, ~1056 | Uchino et al. 2007; Trakunjae et al. 2021 |
| Glycogen | ~1150, ~1082, ~1049, ~1026 | Fabienne Quilès et al. 2012; Holman et al 2009; König et al. 1982 |
| Polyphosphate | ~1275-1260 ($PO_2$ groups) | Zhang et al. 2016 |
| **Osmolytes** | | |
| Trehalose ($C_{12}H_{22}O_{11}$) | ~1150-1000 (~1150, ~1130, ~1100, ~1085, ~1062, ~1031, ~1016, ~1000) (trehalose moieties) | Caccamo et al. 2022; Kuroiwa et al. 2015; Akao et al. 20019 |
| Mannitol ($C_6H_{14}O_6$) | ~1450, ~1420, ~1280, ~1082, ~1019 | Bayón & Rojas, 2017 |
| Ectoine ($C_6H_{10}N_2O_2$) | ~1602, ~1388, ~1308, ~1292, ~1270 | Golovina et al. 2014; Anburajan et al. 20 |
| Proline ($C_5H_9NO_2$) | ~1625, ~1616, ~1565, ~1409, ~1380, | Liu et al. 2021 |
| Glutamate ($C_5H_9NO_4$) | ~1596, ~1562, ~1450, ~1333 | Parikh et al. 2011 |
| Glycine betaine ($C_5H_{11}NO_2$) | ~1493, ~1481, ~1453, ~1400, ~1335, ~1234, ~1122 | Kirschenbaum, 1963 |
| Carnitine ($C_7H_{15}NO_3$) | ~1731, ~1724, ~1685, ~1580, ~1483, ~1411, ~1383, ~968 | Acosta et al. 2020 |

**Supplemental Table 4:** Characteristic key absorption peak frequencies in the mid-infrared spectral range from 1800 to 1000 $cm^{-1}$ for Arthrobacter-SAM under different types of stress. The prominent adsorption peak frequencies for the control cells are provided in black font and the changes under the stress are recorded in the adjacent columns. Where no change is recorded, the peak from the control did not appear under the stress condition. Peak wavenumbers in orange indicate a new peak not observed under control conditions but might appear in one, or more, of the stress conditions. Key: ↑: peak increased in relative intensity under stress. ↓ peak decreased in relative intensity under stress. - the peak was present at the same relative intensity. ↑→ / ↓→: increasing/ decreasing peak intensity accompanied by redshift. ←↑/ ←↓: increasing/ decreasing peak intensity accompanied by a blue shift.

(a) Important carbonyl (C=O) bands of lipids & fatty acids

| **Peak Frequency ($cm^{-1}$)** | **Assignment** | **Prominent adsorption peaks in control & stress experiments** | | | | | |
|---|---|---|---|---|---|---|---|
| | | **Ctl.** | **MS1** | **MS2** | **OS1** | **OS2** | **OS3** |
| 1750 – 1700 | υ carbonyl C=O | 1752 | | ↑ | | | |
| | | 1746 | ↓→ 1742 | ↓ | ↑ | ↓ | ↓ |
| ~1748 – 1740 | υ C=O of ester (in PHA, confirmed by the presence of C-O of alkoxy group at 1311 $cm^{-1}$, and the CH=CH2 at 1027 $cm^{-1}$ (Nwinyi et al., 2017)) | 1740 | | | | ↑ | ↑ |
| | | 1736 | ↑ | ↑ | | ↑ | |
| ~1734 | υ C=O of saturated long-chain esters or ether esters | | | | | | |

(b) Important amide I and amide II bands of proteins

| Peak Frequency (cm$^{-1}$) | Assignment | Prominent adsorption peaks in control & stress experiments | | | | | |
|---|---|---|---|---|---|---|---|
| | | Ctl. | MS1 | MS2 | OS1 | OS2 | OS3 |
| | | 1695 | | | | | ↑ |
| ~1690 – 1610 | υC=O and υC-N of Amide I band components | 1690 | | ↑ | ↑ | ↑ | |
| | | 1683 | | | | | ↑ |
| | | 1676 | ↱ | | | | |
| | | 1659 | ↳ 1657 | ↳ 1657 | ↳ 1657 | ↳ 1657 | ↳ 1655 |
| | | 1639 | ↳ 1638 | ↑ | ↳ 1638 | ↑ | ↓ |
| ~1580 – 1470<br>~1577<br>~1549<br>~1538 | υC-N, υC-C, and δN-H of Amide II band components | 1548 | ↳ 1545 | ↓ | ↑ | ↑ | ↳ 1545 |
| | | 1535 | | | ← 1535 | ↓ | ← 1535 |
| | | 1516 | ↱ 1514 | ↓ | ↱ 1514 | ↴ 1514 | ↱ 1514 |
| ~1518 - 1495 | Aromatic rings (including Tyr & Phe) | 1499 | ↓ | ↳ 1496 | ↳ 1496 | ↳ 1496 | |

(c) Important complex infrared absorption bands of lipids, fatty acids and proteins

| Peak Frequency ($cm^{-1}$) | Assignment | Prominent adsorption peaks in control & stress experiments | | | | | |
|---|---|---|---|---|---|---|---|
| | | Ctl. | MS1 | MS2 | OS1 | OS2 | OS3 |
| ~1475 – 1370<br>~1470<br>~1456<br>~1439 | δC-H of $CH_3$, $CH_2$<br>C-H def of >$CH_2$ | 1467 | ↑ | ↑ | ↓ | | ↑ |
| | | 1456 | ↓ | ↱ 1453 | ↑ | ↳ 1454 | ↴ 1457 |
| | | 1437 | ↓ | ↴ 1441 | ↴ 1440 | | |
| | | 1421 | | | | | |
| | | 1412 | | ↑ | | | |
| ~1440 – 1395 | C-O of carboxylic acid or<br>-COO of carboxylate | 1407 | | | | | ↑ |
| | | 1400 | ↑ | ↑ | ↲ 1407 | ↑ | |
| | | 1389 | — | ↑ | ↑ | ↑ | |
| | | 1389 | | ↑ | | | |
| | | 1382 | | | | | |
| ~1370 +/– 20 | Bending (δ, ρ, ω, τ) of OH of primary & secondary alcohols | 1371 | | ↓ | ↑ | ↑ | ← 1373 |

Table c cont.

| Peak Frequency ($cm^{-1}$) | Assignment | Prominent adsorption peaks in control & stress experiments | | | | | |
|---|---|---|---|---|---|---|---|
| | | Ctl. | MS1 | MS2 | OS1 | OS2 | OS3 |
| ~1310 – 1240 | Amide III components of protein<br>1300 $cm^{-1}$ α-helix<br>1285 & 1245 $cm^{-1}$ random coil<br>1265 & 1235 $cm^{-1}$ β-turn spectrum | 1317 | | | ↑ | | |
| | | 1315 | | ↑ | | | |
| | | 1312 | ↑ | | | ↑ | |
| | | 1308 | | | | | |
| | | 1305 | | | | ↑ | |
| | | 1304 | | | ↑ | | |
| | | 1302 | | | | | ↑ |
| | | 1286 | | ↑ | | ↑ | ↑ |
| | | 1261 | | ↑ | | | |
| | | 1245 | | ↑ | | | |
| | | 1238 | | ↑ | | | |

(d) Important infrared absorption bands of carbohydrates

| Peak Frequency ($cm^{-1}$) | Assignment | Prominent adsorption peaks in control & stress experiments | | | | | |
|---|---|---|---|---|---|---|---|
| | | Ctl. | MS1 | MS2 | OS1 | OS2 | OS3 |
| | | 1253 | | | ↑ (1255) | | ↑ |
| | $\upsilon_{as}$ P-O of $>PO_2^-$ phosphodiester | 1243 | ↑ | | | | |
| 1250 – 1220 | 1210 -1215 $cm^{-1}$ for Z-form<br>1222 -1225 $cm^{-1}$ for B-form | 1238 | ↓ | | ↓ | | |
| | 1240 -1250 $cm^{-1}$ for A-form | 1235 | ↑ | | | | ↑ |
| | | 1223 | | ↑ (1221) | ↑ (1230) | | |
| | | 1218 | | | | | ↑ |
| | δ C-H of $CH_2$ coupled with C-O modes | 1211 | | ↓ | ↓ (1208) | ↑ | |
| 1240 – 1171 | | 1195 | ↓ | | ↓ (1186) | ↓ | |
| | | 1181 | | | | | ↑ |
| | υ of C-O-C linkages - | 1170 | | ↑ | | | |
| | 1160 $cm^{-1}$ carbohydrate rich bands (Nichols, 1985) | 1168 | ↑ | | | ↑ | |
| | | 1160 | ↓ | | ↑ (1166) | | |
| | | 1159 | | ↑ | | | |
| | | 1144 | | | | ↓ | |
| | | 1133 | ↑ | | | | |
| 1140 – 1070 | $\upsilon_{as}$ C-O of saturated unbranched ethers in carbohydrates - | 1125 | ↓ | ↓ (1120) | | | |
| | | 1121 | ↑ | | | | ↑ |
| 1300 – 1200/<br>1050 - 1000 | $\upsilon_{as}$ C-O of mixed (alkyl/ aryl) ethers in carbohydrates - | 1113 | | | ↑ | ↑ | ↑ |
| | | 1110 | | | | | |
| 1200 – 1000 | $\upsilon_s$ C-O, C-C, and C-O-H, C-O-C def of carbohydrates | 1103 | | | ↑ | | ↑ |

(d) Important infrared absorption bands of carbohydrates (cont.)

| Peak Frequency ($cm^{-1}$) | Assignment | Prominent adsorption peaks in control & stress experiments | | | | | |
|---|---|---|---|---|---|---|---|
| | | Ctl. | MS1 | MS2 | OS1 | OS2 | OS3 |
| | | 1099 | | ↑ | ↑ | ↑ | |
| 1090 – 1085 | $\upsilon_{as}$ P-O of >$PO_2^-$ | 1090 | | | | | |
| | | 1083 | ↓ | | ↑ | | ↓ |
| 1150 – 1070 | $\upsilon_{as}$ C-O of secondary hydroxyl - 1150: carbohydrate of bacterial gum (Nichols, 1985) | 1081 | ↑ | ↑ | | ↑ | ↑ |
| | | 1076 | ↓ | | ↓ | | ↓ |
| | | 1069 | | | ↑ | | |
| | | 1061 | ↑→ 1060 | ↑→ 1060 | | | |
| | | 1059 | | | ↑ | | ↑ |
| | | 1054 | ↑ | | | | |
| | | 1045 | ↓ | | | | |
| | | 1040 | ↑ | | | | |
| 1075 – 1000 | $\upsilon_{as}$ C-O of primary hydroxyl - of carbohydrates | 1038 | | ↑ | | | ↑ |
| | | 1034 | ↓ | | | ↓ 1033 | |
| | | 1032 | | ↑ | | | |
| | | 1030 | ↑ | | | | |
| | | 1028 | | | | | ↓ 1029 |
| | | 1024 | | | | | |
| | | 1023 | | ↑ | | | |
| | | 1021 | ↑ | | ↑ | | |
| | | 1018 | | | ↑ | | ↑ |
| | | 1014 | | | | ↑ | |
| | | 1005 | | | | | ↑ |

**Supplemental Table 5:** Characteristic key absorption frequencies in the mid-infrared spectral range from 1800 to 1000 $cm^{-1}$ for Arthrobacter-HTF under different types of stress. Refer to supplemental table 4 for additional information on adsorption peak assignment, and a table key.

(a) Lipid & fatty acid carbonyl (C=O) bonds region (~1780 - 1700 cm-1).

| Peak Frequency ($cm^{-1}$) | Assignment | Prominent adsorption peaks in control & stress experiments | | | | | |
|---|---|---|---|---|---|---|---|
| | | Ctl. | MS1 | MS2 | OS1 | OS2 | OS3 |
| 1750 – 1700 | υ carbonyl C=O | 1750 | | | ↑ | ↑ | |
| | | 1748 | | | | | |
| ~1748 – 1740 | υ C=O of ester (in PHA, | 1745 | ↓ | ↓ | ↑ | | |
| | | 1743 | | | ↑ | | ↑ |
| | | 1738 | | | ↑ | ↑ | |
| | | 1736 | ↑ | ↑ | | | |
| ~1734 | υ C=O of saturated long-chain esters or ether esters | 1733 | | | ↑ | | |
| | | 1731 | ↑ | | | | |
| ~1722 | υ C=O of saturated aldehydes | 1724 | | | | ↑ | ↑ |

(b) Important amide I and amide II bands of proteins

| Peak Frequency ($cm^{-1}$) | Assignment | Prominent adsorption peaks in control & stress experiments | | | | | |
|---|---|---|---|---|---|---|---|
| | | Ctl. | MS1 | MS2 | OS1 | OS2 | OS3 |
| 1715 – 1680 | υ-carbonyl C=O of nucleic acids: | 1695 | | | ↱ 1693 | ↴ 1699 | ↱ 1691 |
| | | 1688 | | ↑ | | | |
| ~1690 – 1610 | υC=O and υC-N of | 1682 | | | ↑ | | |
| | Amide I band components | 1675 | | ↑ | | | ↑ |
| | | 1657 | ↱ 1656 | ↳ 1656 | ↑ | ↓ | ↴ 1658 |
| | | 1640 | ↓ | ↑ | ↓ | ↓ | — |
| | | 1635 | ↓ | | | | |
| | | 1616 | | | | ↑ | |
| | | 1577 | ↑ | | | ↓ | |
| | | 1569 | | | | | ↑ |
| | | 1560 | | | | ↑ | |
| ~1580 – 1470 | υC-N, υ C-C, and δ N-H of | 1558 | | | | | |
| ~1577 | Amide II band components | 1549 | ↓ | | ↓ | | ↑ |
| ~1549 | | 1541 | | | | ↑ | |
| ~1538 | | 1537 | ↑ | ↱ 1553 | ↱ 1553 | ↓ | |
| ~1518 - 1495 | | 1527 | | | | | ↑ |
| | Aromatic rings | 1516 | ↲ 1520 | ↱ 1514 | ↲ 1520 | ↱ 1514 | ↑ |
| ~1520 – 1490 | (Incl. Tyr and Phe) | 1495 | ↑ | ↑ | | ↑ | |

(c) Important complex infrared absorption bands of lipids, fatty acids and proteins

| Peak Frequency ($cm^{-1}$) | Assignment | Prominent adsorption peaks in control & stress experiments | | | | | |
|---|---|---|---|---|---|---|---|
| | | Ctl. | MS1 | MS2 | OS1 | OS2 | OS3 |
| ~1475 – 1370<br>~1470<br>~1456<br>~1439 | δC-H of $CH_3$, $CH_2$<br>C-H def of >$CH_2$ | 1470 | — | — | ↓ | ↑ 1469 | ↑ 1469 |
| | | 1456 | ↑ 1457 | ↓ | ↓ | | ↓ |
| | | 1440 | ↑ | | | ↑ | |
| | | 1438 | | ↑ | | | ↑ |
| | | 1436 | | | ↑ | | |
| | | 1426 | | | | ↑ | ↑ |
| ~1440 – 1395 | C-O of carboxylic acid or<br>-COO of carboxylate | 1418 | ↑ 1420 | | ↓ | | |
| | | 1426 | | | ↑ | ↑ | ↑ |
| | | 1403 | | ↑ | | ↑ | ↑ |
| | | 1418 | | | | | |
| | | 1393 | ↑ | | ↑ | ↑ | ↑ |
| | | 1388 | | ↓ 1390 | | | |
| ~1370 +/– 20 | Bending (δ, ρ, ω, τ) of OH of primary & secondary alcohols (1370 cm-1 indicate carbohydrate-rich bands ) | 1374 | ↑ 1378 | ↑ 1373 | ↓ | ↑ | |
| | | 1354 | | ↑ | ↑ | ↑ | |
| | | 1347 | | | | | ↑ |
| | | 1340 | | | ↓ | | |
| | | 1332 | | ↑ | ↑ | ↑ | ↑ |
| | | 1311 | — | ↓ 1315 | ↑ 1317 | | |
| | | 1306 | | | | ↑ | |
| | | 1300 | | | ↑ | | |
| | | 1297 | ↑ | | | | ↑ |
| | | 1287 | | ↑ | | | |
| ~1310 – 1240 | Amide III components of protein<br>1300 $cm^{-1}$ α-helix<br>1285 & 1245 $cm^{-1}$ random coil<br>1265 & 1235 $cm^{-1}$ β-turn spectrum | 1285 | | | | ↑ 1287 | |
| | | 1281 | | | | | ↑ |
| | | 1274 | | | ↑ | | |
| | | 1264 | | | ↑ | | |
| | | 1252 | ↑ | | | ↑ | |
| | | 1251 | | | | | ↑ |
| | | 1241 | | ↑ 1246 | ↑ 1240 | ↑ 1246 | |

(d) Important infrared absorption bands of carbohydrates

| Peak Frequency ($cm^{-1}$) | Assignment | Prominent adsorption peaks in control & stress experiments | | | | | |
|---|---|---|---|---|---|---|---|
| | | Ctl. | MS1 | MS2 | OS1 | OS2 | OS3 |
| 1200 – 1000 | $υ_s$ C-O, C-C, and C-O-H, C-O-C def of carbohydrates | 1232 | | ↑ | ↑ (1238) | | ↑ |
| 1300 – 1200/ 1050 - 1000 | $υ_{as}$ C-O of mixed (alkyl/ aryl) ethers in carbohydrates - 1240: carbohydrate rich bands | 1222 | | ↓ (1218) | ↓ (1225) | | ↑ (1220) |
| | | 1217 | | | ↑ | ↑ | |
| 1250 – 1220 | $υ_{as}$ P-O of >$PO_2^-$ phosphodiester | 1208 | | | | | |
| | | 1201 | | | ↑ | | ↑ |
| | | 1187 | | | ↑ | ↑ | ↑ |
| | | 1179 | | | ↑ | | |
| | δ C-H of $CH_2$ coupled with C-O modes | 1174 | ↑ | | | | |
| 1240 – 1171 | | 1168 | | ↑ | | ↑ | |
| | υ of C-O-C linkages - 1160: carbohydrate rich bands (Nichols, 1985) | 1165 | ↑ | | | | ↑ |
| | | 1159 | | | ↑ | | ↑ |
| | | 1155 | ↑ | | | | |
| | | 1154 | | | | ↑ | |
| | | 1146 | | | ↑ | | ↑ |
| | | 1132 | | | ↓ | ↑ | |
| | | 1126 | | | | | ↑ |
| | | 1122 | ↑ | ↑ | | | |
| 1140 – 1070 | $υ_{as}$ C-O of saturated unbranched ethers in carbohydrates - | 1114 | ↑ | ↑ | | | |
| | | 1110 | | | | | ↑ |
| | | 1104 | ↑ | ↑ | | | |
| 1090 – 1085 | $υ_{as}$ P-O of >$PO_2^-$ | 1087 | | | ↑ | ↓ | |
| 1150 – 1070 | $υ_{as}$ C-O of secondary hydroxyl - 1150: carbohydrate of bacterial gum (Nichols, 1985) | 1080 | | | | | ↑ |
| | | 1077 | ↓ | ↑ | ↓ | | |
| | | 1075 | | | | ↓ | ↓ |

(d) Important infrared absorption bands of carbohydrates

| Peak Frequency ($cm^{-1}$) | Assignment | Prominent adsorption peaks in control & stress experiments | | | | | |
|---|---|---|---|---|---|---|---|
| | | Ctl. | MS1 | MS2 | OS1 | OS2 | OS3 |
| 1075 – 1000 | $\upsilon_{as}$ C-O of primary hydroxyl - of carbohydrates | 1067 | ↑ | | | | |
| | | 1062 | | ↑ | | ↓ | ↑→ 1061 |
| | | 1055 | | | ↑ | ↑ | |
| | | 1037 | | | | | |
| | | 1032 | ↑ | | | | ↑ |
| | | 1030 | ↑ | | | | |
| | | 1028 | | | | | |
| | | 1021 | | ↑ | ↑ | ↑ | ↑ |
| | | 1011 | ↑ | | ↑ | ↑ | |
| | | 1005 | | | | | ↑ |

**Supplemental Table 6:** Characteristic key absorption frequencies in the mid-infrared spectral range from 1800 to 1000 $cm^{-1}$ for Bacillus-SAM under different types of stress. Refer to supplemental table 4 for additional information on adsorption peak assignment, and a table key.

(a) Important carbonyl (C=O) bands of lipids & fatty acids

| Peak Frequency ($cm^{-1}$) | Assignment |
|---|---|
| 1750 – 1700 | υ carbonyl C=O |
| ~1748 – 1740 | υ C=O of ester (in PHA) |
| ~1734 | υ C=O of saturated long-chain esters or ether esters |
| ~1722 | υ C=O of saturated aldehydes |
| ~1715 | υ C=O of saturated aliphatic ketones |
| 1713 - 1709 | υ C=O of saturated aliphatic carboxylic acids |

| Prominent adsorption peaks in control & stress experiments | | | | | |
|---|---|---|---|---|---|
| Ctl. | MS1 | MS2 | OS1 | OS2 | OS3 |
| 1748 | — | ↓ 1742 | ↓ | ↑ | ↓ |
| 1745 | ↑ | ↓ | | ↓ | |
| 1741 | | | | ↑ | ↑ |
| 1735 | | | ↑ | | |
| 1725 | ↑ | ↓ | ↓ | ↓ | ↓ |
| 1722 | | | | ↑ | |
| 1711 | | | ↑ | | ↑ |
| 1700 | | | | ↑ | |

(b Important amide I and amide II bands of proteins

| Peak Frequency (cm$^{-1}$) | Assignment | Prominent adsorption peaks in control & stress experiments | | | | | |
|---|---|---|---|---|---|---|---|
| | | Ctl. | MS1 | MS2 | OS1 | OS2 | OS3 |
| 1715 – 1680 | υ-carbonyl C=O of nucleic acids: | 1690 | | | ↑ | ↑ | |
| | | 1669 | | | | ↑ | |
| | | 1660 | ↑ | ↑→ 1659 | ←↑ 1665 | ↑→ 1657 | |
| | | 1658 | | | | | ↑ |
| | | 1650 | | | | | ↑ |
| | | 1640 | | ↑ | | | |
| ~1690 – 1610 | υC=O and υC-N of Amide I band components | 1638 | | | | ↑ | ↑ |
| | | 1632 | | | | | ↑ |
| | | 1629 | ↓ | | | | ↓ |
| | | 1594 | | | ↑ | | |
| | | 1578 | | | ↑ | ↑ | |
| | | 1560 | | | ↑ | ↓ | ↓ |
| ~1580 – 1470<br>~1577 | υC-N, υ C-C, and δ N-H of Amide II band components | 1543 | ↑ | ↓ | ↑→ 1541 | ↑→ 1541 | |
| ~1549 | | 1521 | | | | ↑ | |
| ~1538 | | 1517 | ↑ | ↓ | ←↑ 1619 | | ↑ |
| ~1518 - 1495 | | 1511 | | | | ↑ | |
| ~1520 – 1490 | Aromatic rings (Incl. Tyr and Phe) | 1498 | | | | ↑ | ↓ |

(c) Important complex infrared absorption bands of lipids, fatty acids and proteins

| Peak Frequency ($cm^{-1}$) | Assignment | Prominent adsorption peaks in control & stress experiments | | | | | |
|---|---|---|---|---|---|---|---|
| | | Ctl. | MS1 | MS2 | OS1 | OS2 | OS3 |
| ~1475 – 1370 | | 1473 | | | | ↑ | ↓ |
| ~1470 | | 1468 | ↓ | ↑ | | ↑ | ↑ |
| ~1456 | δC-H of $CH_3$, $CH_2$ | 1456 | ↑ | ↓ | ↓ | ↑ | ↑ |
| ~1439 | C-H def of >$CH_2$ | 1440 | ↑ | | | | |
| | | 1425 | | | ↑ | ↑ | |
| ~1440 – 1395 | C-O of carboxylic acid or -COO of carboxylate | 1417 | | | | | — |
| | | 1413 | | | | ↑ | |
| | Bending (δ, ρ, ω, τ) of OH of primary & secondary alcohols (1370 cm-1 indicate carbohydrate-rich bands ) | 1381 | ↑ | ↓ | ↑ | ↓ | ↑ |
| ~1370 +/– 20 | | 1371 | | | ↑ | ↑ | ↓ |
| | | 1357 | | | | ↑ | |
| | | 1348 | | | | ↑ | |
| | | 1346 | | | ↑ | | ↓ |
| | | 1338 | | | | ↑ | |
| | | 1332 | | | ↑ | | ↓ |
| | | 1327 | | | | ↑ | |
| | | 1314 | | | | ↑ | |
| | | 1309 | → 1311 | → 1307 | — | | ↓ |
| | Amide III components of protein<br>1300 $cm^{-1}$ α-helix<br>1285 & 1245 $cm^{-1}$ random coil<br>1265 & 1235 $cm^{-1}$ β-turn spectrum | 1293 | → 1292 | ↓ | ↑ | | ↓ |
| ~1310 – 1240 | | 1278 | ↑ | | | | |
| | | 1265 | | | ↓ | ↑ | |
| | | 1262 | ↓ | ↓ | | | ↓ |
| | | 1255 | | | ↓ | ↑ | |
| | | 1246 | | | | ↑ | ↓ |

(d) Important infrared absorption bands of carbohydrates

| Peak Frequency ($cm^{-1}$) | Assignment | Prominent adsorption peaks in control & stress experiments | | | | | |
|---|---|---|---|---|---|---|---|
| | | Ctl. | MS1 | MS2 | OS1 | OS2 | OS3 |
| 1250 – 1220 | $υ_{as}$ P-O of $>PO_2^-$ phosphodiester | 1230 | ↑ | ↓ | | ↓ | — |
| | | 1222 | | | ↑ | | |
| | δ C-H of $CH_2$ coupled with C-O modes | 1209 | | | | ↑ | |
| 1240 – 1171 | | 1203 | | | ↑ | | |
| | υ of C-O-C linkages - 1160: carbohydrate rich bands | 1194 | | | | ↑ | |
| | | 1183 | ↑ | | | | ↑ |
| | | 1177 | | | | ↑ | |
| | | 1172 | | ↑ | ↑ | | ↓ |
| | | 1163 | | | | ↑ | |
| | | 1154 | ↑ | | ↑ | ↑ | ↓ |
| | | 1142 | | | | ↑ | |
| 1140 – 1070 | $υ_{as}$ C-O of saturated unbranched ethers in carbohydrates - | 1138 | | | ↑ | | |
| | | 1133 | ↑ | ↓ | | ↑ | ↑ |
| | | 1129 | | | ↑ | | |
| 1200 – 1000 | $υ_s$ C-O, C-C, and C-O-H, C-O-C def of carbohydrates | | | | | | |
| 1300 – 1200/ 1050 - 1000 | $υ_{as}$ C-O of mixed (alkyl/ aryl) ethers in carbohydrates - 1240: carbohydrate rich bands | | | | | | |

(d) Important infrared absorption bands of carbohydrates

| Peak Frequency ($cm^{-1}$) | Assignment | Prominent adsorption peaks in control & stress experiments | | | | | |
|---|---|---|---|---|---|---|---|
| | | Ctl. | MS1 | MS2 | OS1 | OS2 | OS3 |
| | | 1090 | | | | ↑ | |
| | | 1083 | | | ↑ | | |
| 1150 – 1070 | $\upsilon_{as}$ C-O of secondary hydroxyl - 1150: carbohydrate of bacterial gum | 1080 | | ↑ | | | |
| | | 1073 | | | | ↑ | |
| | | 1069 | | | ↑ | | |
| 1090 – 1085 | $\upsilon_{as}$ P-O of $>PO_2^-$ | 1059 | | ↓ | ↑ | | ↓ |
| | | 1043 | ↓ | ↑ | | | — |
| | | 1035 | | | ↑ | ↑ | |
| 1075 – 1000 | $\upsilon_{as}$ C-O of primary hydroxyl - of carbohydrates | 1030 | ↑ | | ↑ | | ↑ |
| | | 1025 | | | | ↑ | |
| | | 1013 | ↑ | ↑ | ↑ | | ↑ |

**Supplemental Table 7:** Characteristic key absorption frequencies in the mid-infrared spectral range from 1800 to 1000 $cm^{-1}$ for Bacillus-HTF under different types of stress. Refer to supplemental table 4 for additional information on adsorption peak assignment, and a table key.

(a) Important carbonyl (C=O) bands of lipids & fatty acids

| Peak Frequency ($cm^{-1}$) | Assignment |
|---|---|
| 1750 – 1700 | υ carbonyl C=O |
| ~1748 – 1740 | υ C=O of ester |
| ~1734 | υ C=O of saturated long-chain esters or ether esters |
| ~1722 | υ C=O of saturated aldehydes |
| ~1715 | υ C=O of saturated aliphatic ketones |
| 1713 - 1709 | υ C=O of saturated aliphatic carboxylic acids |

| Prominent adsorption peaks in control & stress experiments | | | | | |
|---|---|---|---|---|---|
| Ctl. | MS1 | MS2 | OS1 | OS2 | OS3 |
| 1749 | | | ↓ | ↑ | |
| 1747 | ↓ | ↓ | | ↓ | ↓ |
| 1743 | | ↑ | | | ↑ |
| 1737 | ↑ | | ↑ | ↑ | |
| 1733 | | ↑ | ↓ | | |
| 1731 | ↓ | ↓ | | | ↓ |
| 1716 | | | | | ↑ |
| 1714 | ↓ | ↓ | ↓ | | ↓ |
| 1702 | | | ↑ | | |

(b) Important amide I and amide II bands of proteins

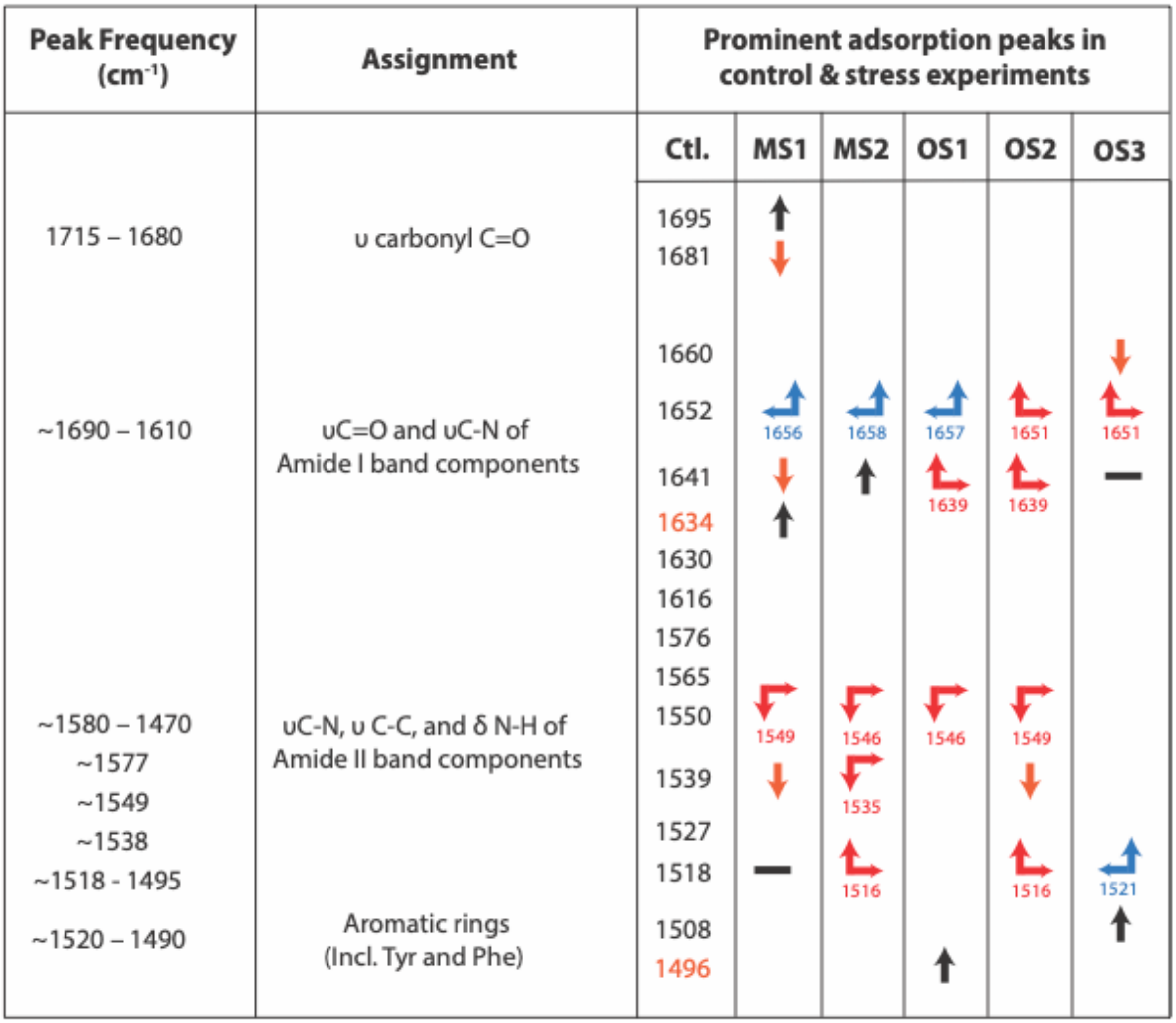

| Peak Frequency ($cm^{-1}$) | Assignment | Prominent adsorption peaks in control & stress experiments | | | | | |
|---|---|---|---|---|---|---|---|
| | | Ctl. | MS1 | MS2 | OS1 | OS2 | OS3 |
| 1715 – 1680 | υ carbonyl C=O | 1695 | ↑ | | | | |
| | | 1681 | ↓ | | | | |
| | | 1660 | | | | | ↓ |
| ~1690 – 1610 | υC=O and υC-N of Amide I band components | 1652 | 1656 | 1658 | 1657 | 1651 | 1651 |
| | | 1641 | ↓ | ↑ | 1639 | 1639 | — |
| | | 1634 | ↑ | | | | |
| | | 1630 | | | | | |
| | | 1616 | | | | | |
| | | 1576 | | | | | |
| | | 1565 | | | | | |
| ~1580 – 1470 | υC-N, υ C-C, and δ N-H of Amide II band components | 1550 | 1549 | 1546 | 1546 | 1549 | |
| ~1577 | | | | | | | |
| ~1549 | | 1539 | ↓ | 1535 | | ↓ | |
| ~1538 | | 1527 | | | | | |
| ~1518 - 1495 | | 1518 | — | 1516 | | 1516 | 1521 |
| ~1520 – 1490 | Aromatic rings (Incl. Tyr and Phe) | 1508 | | | | | ↑ |
| | | 1496 | | | ↑ | | |

(c) Important complex infrared absorption bands of lipids, fatty acids and proteins

| Peak Frequency ($cm^{-1}$) | Assignment | Prominent adsorption peaks in control & stress experiments | | | | | |
|---|---|---|---|---|---|---|---|
| | | Ctl. | MS1 | MS2 | OS1 | OS2 | OS3 |
| | | 1469 | ↑ | ↑ | — | ↑ | ↑ |
| ~1475 – 1370 | | 1453 | ↳ 1456 | ↳ 1456 | ↳ 1456 | ↓ | ↓ |
| ~1470 | δC-H of $CH_3$, $CH_2$ | 1442 | | | | | ↑ |
| ~1456 | C-H def of >$CH_2$ | 1438 | ↑ | | ↑ | ↑ | |
| ~1439 | | 1433 | | | | | ↑ |
| ~1440 – 1395 | C-O of carboxylic acid or -COO of carboxylate | 1430 | | | | | |
| | | 1426 | | | | | |
| | | 1421 | ↑ | | ↑ | | |
| | | 1414 | | | | ↓ | ↓ |
| | | 1401 | | | | | ↑ |
| | | 1398 | | — | — | | ↓ |
| | | 1395 | | ↑ | | | |
| | | 1385 | | ↓ | ↳ 1389 | ↓ | ↑ |
| ~1370 +/– 20 | Bending (δ, ρ, ω, τ) of OH of primary & secondary alcohols | 1374 | | ↳ 1372 | | ↑ | ↑ |
| | | 1368 | | ↑ | | | |
| | | 1363 | | | | | |
| | | 1340 | | ↓ | ↳ 1343 | | ↓ |
| | | 1334 | | | | ↑ | |
| | | 1323 | | ↑ | | | |
| | | 1317 | ↑ | ↑ | | ↑ | ↳ 1319 |
| ~1310 – 1240 | Amide III components of protein<br>1300 $cm^{-1}$ α-helix<br>1285 & 1245 $cm^{-1}$ random coil<br>1265 & 1235 $cm^{-1}$ β-turn spectrum | 1312 | | ↑ | | | |
| | | 1309 | | | | ↑ | |
| | | 1295 | | ↓ | — | ↑ | — |
| | | 1288 | ↑ | | | | |
| | | 1283 | | | | ↑ | |
| | | 1277 | | ↑ | ↑ | | |
| | | 1268 | | | | — | — |
| | | 1251 | ↳ 1248 | ↳ 1250 | ↳ 1247 | ↳ 1256 | ↳ 1250 |

(d) Important infrared absorption bands of carbohydrates

| Peak Frequency ($cm^{-1}$) | Assignment | Prominent adsorption peaks in control & stress experiments | | | | | |
|---|---|---|---|---|---|---|---|
| | | Ctl. | MS1 | MS2 | OS1 | OS2 | OS3 |
| 1250 – 1220 | $\upsilon_{as}$ P-O of >$PO_2^-$ phosphodiester 1210 -1215 $cm^{-1}$ for Z-form 1222 -1225 $cm^{-1}$ for B-form 1240 -1250 $cm^{-1}$ for A-form | 1232 | | ↓ | ↓ | | ↑ |
| | | 1225 | | ↓ | ↓ | ↱ 1222 | |
| | | 1218 | | | ↑ | | ↑ |
| | | 1214 | ↑ | ↑ | | | |
| | | 1204 | | | | ↓ | ↑ |
| | | 1201 | | | ↑ | | |
| 1240 – 1171 | δ C-H of $CH_2$ coupled with C-O modes υ of C-O-C linkages - 1160: carbohydrate rich bands (Nichols, 1985) | 1198 | ↑ | ↑ | | | |
| | | 1187 | | | ↳ 1189 | | |
| | | 1179 | | | ↱ 1177 | | |
| | | 1169 | ↳ 1170 | ↱ 1167 | ↳ 1170 | ↳ 1172 | ↲ 1173 |
| | | 1160 | | | | ↑ | ↑ |
| | | 1158 | ↑ | ↑ | ↑ | | |
| | | 1153 | | | | ↑ | ↑ |
| | | 1147 | ↓ | ↓ | | | |
| | | 1137 | | | | ↑ | ↑ |
| | | 1127 | ↑ | | | | |
| 1140 – 1070 | $\upsilon_{as}$ C-O of saturated unbranched ethers in carbohydrates - | 1124 | | | | | |
| | | 1122 | | ↑ | ↑ | | ↑ |
| | | 1115 | | | | | |
| 1300 – 1200/ 1050 - 1000 | $\upsilon_{as}$ C-O of mixed (alkyl/ aryl) ethers in carbohydrates - 1240: carbohydrate rich bands (Nichols, 1985) | 1110 | ↑ | | ↑ | | |
| | | 1102 | | ↑ | | | ↑ |
| | | 1093 | | | ↑ | | |
| 1200 – 1000 | $\upsilon_s$ C-O, C-C, and C-O-H, C-O-C def of carbohydrates | | | | | | |

(d) Important infrared absorption bands of carbohydrates (cont.)

| Peak Frequency ($cm^{-1}$) | Assignment | Prominent adsorption peaks in control & stress experiments | | | | | |
|---|---|---|---|---|---|---|---|
| | | Ctl. | MS1 | MS2 | OS1 | OS2 | OS3 |
| | | 1081 | ↓ 1086 | | ↑ | ↓ | ↑ |
| 1150 – 1070 | $\upsilon_{as}$ C-O of secondary hydroxyl - 1150: carbohydrate of bacterial gum (Nichols, 1985) | 1077 | ↑ | | | | |
| | | 1068 | ↓ 1067 | | | ↓ | ↓ |
| | | 1059 | ↑ | | | | |
| 1090 – 1085 | $\upsilon_{as}$ P-O of $>PO_2^-$ | 1056 | | | ↑ | ↑ | |
| | | 1044 | | | ↑ | | ↑ |
| | | 1032 | ↑ | ↑ | | | |
| | | 1028 | | | | ↓ | ↑ |
| 1075 – 1000 | $\upsilon_{as}$ C-O of primary hydroxyl - of carbohydrates | 1024 | | ↑ | | | |
| | | 1017 | ↑ | | | | |
| | | 1006 | | | ↑ | | |

**Supplemental Table 8:** Characteristic key absorption frequencies in the mid-infrared spectral range from 1800 to 1000 cm$^{-1}$ for Enterobacter-SAM under different types of stress. Refer to supplemental table 3 for additional information on adsorption peak assignment, and a table key.

(a) Important carbonyl (C=O) bands of lipids & fatty acids .

| Peak Frequency (cm$^{-1}$) | Assignment | Prominent adsorption peaks in control & stress experiments | | | | | |
|---|---|---|---|---|---|---|---|
| | | Ctl. | MS1 | MS2 | OS1 | OS2 | OS3 |
| 1750 – 1700 | υ carbonyl C=O | 1749 | | ↑ (1750) | ↑ | | |
| ~1748 – 1740 | υ C=O of ester (in PHA, confirmed by the presence of C-O of alkoxy group at 1311 cm$^{-1}$, and the CH=CH2 at 1027 cm$^{-1}$ (Nwinyi et al., 2017)) | 1744 | | | | ↑ | ↑ |
| | | 1740 | | ↑ | | | |
| | | 1737 | ↑ | ↑ (1740) | | | ↑ |
| | | 1736.8 | | | | ↓ | |
| ~1734 | υ C=O of saturated long-chain esters or ether esters | 1736 | ↓ | ↓ | ↑ (1335) | | ↓ |
| | | 1730 | | | | ↑ | |
| ~1722 | υ C=O of saturated aldehydes | 1721 | ↑ | | | | |
| ~1715 | υ C=O of saturated aliphatic ketones | 1720 | ↓ | | ↑ | ↓ | ↓ |
| | | 1714 | | | | ↑ | |
| 1713 - 1709 | υ C=O of saturated aliphatic carboxylic acids | 1702 | ↑ | | | | |

(b) Important amide I and amide II bands of proteins

| Peak Frequency ($cm^{-1}$) | Assignment |
|---|---|
| 1715 – 1680 | υ carbonyl C=O |
| ~1690 – 1610 | υC=O and υC-N of Amide I band components |
| ~1580 – 1470<br>~1577<br>~1549<br>~1538<br>~1518 - 1495 | υC-N, υ C-C, and δ N-H of Amide II band components |
| ~1520 – 1490 | Aromatic rings (Incl. Tyr and Phe) |

| Prominent adsorption peaks in control & stress experiments | | | | | |
|---|---|---|---|---|---|
| Ctl. | MS1 | MS2 | OS1 | OS2 | OS3 |
| 1696 | | ↑ | | ↑ | |
| 1694 | | | ↑ | | |
| 1690 | ↑ | | | | |
| 1688.8 | | | | | ↑ |
| 1683 | | | | ↓ | |
| 1675 | ↑ | | | | |
| 1657.2 | | ↓ | 1655 | 1658 | 1658 |
| 1652 | | | | | ↑ |
| 1639 | 1640 | 1640 | 1638.8 | 1640 | ↓ |
| 1599.5 | | | | ↑ | |
| 1576 | | | ↑ | ↑ | |
| 1570 | | ↑ | | | |
| 1561.5 | ↑ | | | | |
| 1546 | ↑ | 1548 | 1539 | 1545 | 1548 |
| 1540 | | | | | ↑ |
| 1537.6 | 1534.9 | 1541 | | 1535 | |
| 1523 | ↑ | | 1526 | | |
| 1514 | | ↑ | ↑ | ↑ | ↑ |
| 1501 | | | | | |

(c) Important complex infrared absorption bands of lipids, fatty acids and proteins

| Peak Frequency ($cm^{-1}$) | Assignment | Prominent adsorption peaks in control & stress experiments | | | | | |
|---|---|---|---|---|---|---|---|
| | | Ctl. | MS1 | MS2 | OS1 | OS2 | OS3 |
| | | 1469 | 1473 | 1469.8 | ↑ | ↑ | |
| ~1475 – 1370 | | 1459 | | | | | ↑ |
| ~1470 | δC-H of $CH_3$, $CH_2$ | 1457 | ↑ | | ↑ | | |
| ~1456 | C-H def of >$CH_2$ | 1454 | | 1452 | | 1453 | ↑ |
| ~1439 | | 1442 | | | | | ↑ |
| ~1440 – 1395 | C-O of carboxylic acid or -COO of carboxylate | 1440 | 1438 | ↑ | ↓ | ↑ | |
| | | 1422 | 1419 | 1419 | 1414 | 1419 | 1413.7 |
| | | 1403 | 1401.7 | | | ↓ | 1404 |
| | | 1390 | 1387 | 1391 | | | |
| | | 1378 | | 1373 | 1381 | 1376 | |
| ~1370 +/– 20 | Bending (δ, ρ, ω, τ) of OH of primary & secondary alcohols (1370 cm-1 indicate carbohydrate-rich bands ) | 1370 | | | 1374 | | ↑ |
| | | 1357 | | | ↑ | | |
| | | 1353 | | | | | ↑ |
| | | 1348 | ↑ | | | | |
| | | 1345 | | ↑ | | | |
| | | 1341 | | | | | ↑ |
| | | 1338.7 | | | | | |
| | | 1336 | – 1334 | 1334 | 1335 | ↑ | |
| | | 1331 | | | | ↑ | |
| | | 1325 | | | ↑ | | |
| | | 1319 | 1318 | | | | ↑ |
| | | 1312 | | 1315 | 1311 | | |
| | | 1305 | ↑ | | | | ↑ |
| | | 1299 | 1291 | 1297 | 1297.8 | | |
| ~1310 – 1240 | Amide III components of protein<br>1300 $cm^{-1}$ α-helix<br>1285 & 1245 $cm^{-1}$ random coil<br>1265 & 1235 $cm^{-1}$ β-turn spectrum | 1288.5 | | | | | ↑ |
| | | 1280 | 1278 | | ↓ | | |
| | | 1274 | | | | 1273 | ↑ |
| | | 1270.6 | | ↑ | | | |
| | | 1260 | | | ↑ | ↑ | ↑ |
| | | 1253 | | | | | |
| | | 1252 | | | | | ↑ |
| | | 1246 | 1249 | ↑ | ↑ | ↑ | |

(d) Important infrared absorption bands of carbohydrates

| Peak Frequency ($cm^{-1}$) | Assignment | Prominent adsorption peaks in control & stress experiments | | | | | |
|---|---|---|---|---|---|---|---|
| | | Ctl. | MS1 | MS2 | OS1 | OS2 | OS3 |
| 1250 – 1220 | $\upsilon_{as}$ P-O of >$PO_2^-$ phosphodiester | 1234 | 1238 | ↑ | 1238 | 1230 | 1238 |
| | | 1222 | | ↓ | ↑ | 1219 | 1220.1 |
| | | 1209 | ↑ | | ↑ | | |
| 1240 – 1171 | δ C-H of $CH_2$ coupled with C-O modes<br>υ of C-O-C linkages - 1160: carbohydrate rich bands (Nichols, 1985) | 1203 | | 1206.6 | | | 1207 |
| | | 1195 | ↑ | | | | |
| | | 1190 | | | | ↑ | ↑ |
| | | 1183.9 | ↑ | | | | |
| | | 1176 | | | | ↑ | ↑ |
| 1200 – 1000 | $\upsilon_s$ C-O, C-C, and C-O-H, C-O-C def of carbohydrates | 1174 | ↑ | | | | |
| | | 1170 | | | | ↑ | |
| 1300 – 1200/ 1050 - 1000 | $\upsilon_{as}$ C-O of mixed (alkyl/ aryl) ethers in carbohydrates - 1240: carbohydrate rich bands (Nichols, 1985) | 1164 | 1158 | 1163 | 1169 | 1160 | 1169 |
| | | 1152 | | ↑ | | | |

(d) Important infrared absorption bands of carbohydrates (cont.)

| Peak Frequency ($cm^{-1}$) | Assignment | Prominent adsorption peaks in control & stress experiments | | | | | |
|---|---|---|---|---|---|---|---|
| | | Ctl. | MS1 | MS2 | OS1 | OS2 | OS3 |
| | | 1148 | ↑ | | | | |
| | | 1143 | | | | | ↑ |
| | | 1141 | | | | ↓ | |
| | | 1130 | ↑ | | | | |
| 1150 – 1070 | $\upsilon_{as}$ C-O of secondary hydroxyl - 1150: carbohydrate of bacterial gum (Nichols, 1985) | 1119 | ↓ | | 1120 | 1121 | 1117 |
| | | 1110 | | | ↑ | | |
| | | 1104 | | ↑ | | 1106 | |
| | | 1098 | | | ↑ | | |
| 1090 – 1085 | $\upsilon_{as}$ P-O of >$PO_2^-$ | 1091 | 1087 | 1090 | | ↓ | |
| | | 1083 | | ↑ | | | |
| | | 1077 | 1079 | 1080 | | 1076 | 1082 |
| | | 1070 | 1068 | 1069 | 1066 | | ↓ |
| | | 1063 | | | | 1062 | |
| | | 1055 | | ↑ | | | |
| 1075 – 1000 | $\upsilon_{as}$ C-O of primary hydroxyl - of carbohydrates | 1050 | | | | ↑ | ↑ |
| | | 1043 | 1041 | | 1043.8 | | |
| | | 1039 | | | | 1038 | |
| | | 1033 | | ↑ | | | |
| | | 1022 | 1024 | 1020 | 1027 | 1020 | 1025 |
| | | 1016 | | | ↑ | | ↑ |
| | | 1004 | | | | | ↑ |

**Supplemental Table 9:** Characteristic key absorption frequencies in the mid-infrared spectral range from 1800 to 1000 $cm^{-1}$ for Enterobacter_HTF under different types of stress. Refer to supplemental table 4 for additional information on adsorption peak assignment, and a table key.

(a) Important carbonyl (C=O) bands of lipids & fatty acids .

| Peak Frequency ($cm^{-1}$) | Assignment | Prominent adsorption peaks in control & stress experiments | | | | | |
|---|---|---|---|---|---|---|---|
| | | Ctl. | MS1 | MS2 | OS1 | OS2 | OS3 |
| 1750 – 1700 | υ carbonyl C=O | | | | | | |
| | | 1748 | | | 1745 | 1743 | ↑ |
| ~1748 – 1740 | υ C=O of ester (in PHA, confirmed by the presence of C-O of alkoxy group at 1311 $cm^{-1}$, and the CH=CH2 at 1027 $cm^{-1}$ (Nwinyi et al., 2017)) | 1735 | 1737 | 1740 | ↑ | 1734 | 1736 |
| ~1734 | υ C=O of saturated long-chain esters or ether esters | 1722 | ↑ | | | | |
| ~1722 | υ C=O of saturated aldehydes | 1721 | 1717 | | ↑ | | |
| ~1715 | υ C=O of saturated aliphatic ketones | 1716 | ↓ | ↓ | ↑ | ↑ | |
| 1713 - 1709 | υ C=O of saturated aliphatic carboxylic acids | | | | | | |

(b) Important amide I and amide II bands of proteins

| Peak Frequency ($cm^{-1}$) | Assignment | Prominent adsorption peaks in control & stress experiments | | | | | |
|---|---|---|---|---|---|---|---|
| | | Ctl. | MS1 | MS2 | OS1 | OS2 | OS3 |
| 1715 – 1680 | υ-carbonyl C=O | 1696 | 1689.9 | 1691 | 1693.7 | 1695 | 1691 |
| | | 1682 | | | | ↑ | |
| ~1690 – 1610 | υC=O and υC-N of Amide I band components | 1678.7 | | | 1683 | | 1680 |
| | | 1667 | | | ↓ | | |
| | | 1657.7 | 1656.8 | 1658.3 | ↓ | 1656.6 | 1656 |
| | | 1637.6 | 1639 | | 1646 | ↓ | 1638.7 |
| | | 1621 | | | ↑ | | |
| | | 1613 | | | ↑ | | |
| | | 1599 | | | ↑ | | |
| | | 1576 | | 1578 | 1583 | | 1573 |
| | | 1570 | | | ↑ | | |
| ~1580 – 1470 | υC-N, υ C-C, and δ N-H of Amide II band components | 1559 | | | ↑ | | |
| ~1577 | | 1552.2 | ↓ | 1553.9 | | ↓ | |
| ~1549 | | | | | | | |
| ~1538 | | 1546 | 1544 | 1541 | ↓ | ↓ | ↓ |
| ~1518 - 1495 | | 1539 | | | | | ↑ |
| ~1520 – 1490 | Aromatic rings (Incl. Tyr and Phe) | 1537 | ↑ | | | | |
| | | 1536.6 | | | | ↑ | |
| | | 1534 | | | ↑ | | |
| | | 1517 | 1515 | – 1520 | – 1523 | 1516 | 1515 |
| | | 1505 | | | ↑ | | |

(c) Important complex infrared absorption bands of lipids, fatty acids and proteins

| Peak Frequency ($cm^{-1}$) | Assignment | Prominent adsorption peaks in control & stress experiments | | | | | |
|---|---|---|---|---|---|---|---|
| | | Ctl. | MS1 | MS2 | OS1 | OS2 | OS3 |
| | | 1468 | ↓ 1470 | ↑ | ↑ | ↓ | ↓ |
| | | 1464 | | | ↑ | | |
| | | 1454 | ↑ 1455 | ↓ 1455 | ↑ 1443 | ↑ 1455 | ↑ |
| ~1475 – 1370<br>~1470<br>~1456<br>~1439 | δC-H of $CH_3$, $CH_2$<br>C-H def of $>CH_2$ | 1437.5 | ↓ 1441 | ↑ 1441 | ↓ 1440 | ↓ 1440 | ↓ 1437 |
| | | 1434 | | | ↓ | | |
| | | 1420 | | | | ↓ | |
| ~1440 – 1395 | C-O of carboxylic acid or -COO of carboxylate | 1418 | | | ↑ | | ↓ |
| | | 1414 | | ↓ | | | |
| | | 1402 | ↓ | ↓ 1398 | ↓ | | ↓ |
| | | 1393 | | | | | ↑ |
| | | 1385 | ↑ 1390 | ↑ 1386.9 | | ↓ 1392 | ↓ 1383 |
| ~1370 +/– 20 | Bending (δ, ρ, ω, τ) of OH of primary & secondary alcohols (1370 cm-1 indicate carbohydrate-rich bands ) | 1379 | | | | ↑ | |
| | | 1364.4 | ↓ 1370 | ↓ 1373 | ↓ 1358 | ↑ ← 1367 | ↓ 1369 |
| | | 1349 | | | | ↑ | |
| | | 1346 | ↑ | | | | |
| | | 1344 | | | ↑ | | |
| | | 1335 | ↑ 1331 | ↑ 1340 | ↑ 1331 | ↑ | ↑ 1340 |
| | | 1322 | | ↑ | | ↑ | |
| | | 1318 | | | | | ↑ |
| | | 1315 | | | ↑ | | |
| | | 1312 | | | | ↑ | |
| | | 1302 | ↑ | ↓ | | | ↑ |
| | | 1292 | | | | – 1297 | |
| ~1310 – 1240 | Amide III components of protein<br>1300 $cm^{-1}$ α-helix<br>1285 & 1245 $cm^{-1}$ random coil<br>1265 & 1235 $cm^{-1}$ β-turn spectrum | 1286.9 | ↓ 1281 | | | ↑ 1279 | ↓ 1288 |
| | | 1268 | ↑ | | ↑ | | |
| | | 1258.1 | | ↓ 1259 | | ↓ 1260 | ↑ 1264 |
| | | 1251 | | | ↑ | ↓ | |
| | | 1248 | | ↑ | | ↑ | |
| | | 1243 | ↑ | | | | |

(d) Important infrared absorption bands of carbohydrates region

| Peak Frequency ($cm^{-1}$) | Assignment | Prominent adsorption peaks in control & stress experiments | | | | | |
|---|---|---|---|---|---|---|---|
| | | Ctl. | MS1 | MS2 | OS1 | OS2 | OS3 |
| 1250 – 1220 | $\upsilon_{as}$ P-O of $>PO_2^-$ phosphodiester | 1236 | | ↑ | ↑ | ↑ | ↑ |
| | | 1227.7 | 1225 | 1226 | 1221 | 1229 | 1224 |
| | | 1216 | 1213 | 1213 | | | |
| 1240 – 1171 | δ C-H of $CH_2$ coupled with C-O modes | 1193 | | 1190 | 1197 | | |
| | υ of C-O-C linkages - 1160: carbohydrate rich bands | 1183 | | | | ↑ | |
| | | 1172.7 | 1176 | 1172 | 1175 | ↓ | ↓ |
| 1200 – 1000 | $\upsilon_s$ C-O, C-C, and C-O-H, C-O-C def of carbohydrates | 1159.5 | 1163 | 1154 | 1155 | 1153 | 1166 |
| 1300 – 1200/ 1050 - 1000 | $\upsilon_{as}$ C-O of mixed (alkyl/ aryl) ethers in carbohydrates - 1240: carbohydrate rich bands | | | | | | |

(d) Important infrared absorption bands of carbohydrates (cont.)

| Peak Frequency ($cm^{-1}$) | Assignment | Prominent adsorption peaks in control & stress experiments | | | | | |
|---|---|---|---|---|---|---|---|
| | | Ctl. | MS1 | MS2 | OS1 | OS2 | OS3 |
| | | 1142.5 | | 1138 | | | 1150 |
| | | 1136 | | | | | ↑ |
| | | 1133 | | | ↓ | | |
| 1150 – 1070 | $\upsilon_{as}$ C-O of secondary hydroxyl - 1150: carbohydrate of bacterial gum (Nichols, 1985) | 1128 | | | 1126 | ↑ | |
| | | 1114 | ↑ | 1110 | 1109 | 1117 | 1118.9 |
| | | 1098.6 | 1101 | ↓ | | 1098 | 1106 |
| 1090 – 1085 | $\upsilon_{as}$ P-O of $>PO_2^-$ | 1084 | 1082 | 1085 | | 1081 | 1082 |
| | | 1078 | | ↑ | ↑ | | |
| | | 1072 | ↑ | | | | |
| | | 1069 | | | | | ↑ |
| | | 1065 | | 1061 | | 1061 | |
| | | 1059 | ↑ | | | | |
| | | 1053 | | | 1051 | 1051 | 1056 |
| 1075 – 1000 | $\upsilon_{as}$ C-O of primary hydroxyl - of carbohydrates | 1038.9 | 1034 | | 1034 | 1031 | |
| | | 1026 | 1018 | — | 1022 | 1016 | |
| | | 1017 | | ↑ | | | |
| | | 1011 | | | 1008 | | 1009 |

**Supplemental Table 10:** Characteristic key absorption frequencies in the mid-infrared spectral range from 1800 to 1000 $cm^{-1}$ for Sphingomonas-SAM under different types of stress. Refer to supplemental table 4 for additional information on adsorption peak assignment, and a table key.

(a) Important carbonyl (C=O) bands of lipids & fatty acids

| Peak Frequency ($cm^{-1}$) | Assignment | Prominent adsorption peaks in control & stress experiments | | | | | |
|---|---|---|---|---|---|---|---|
| | | Ctl. | MS1 | MS2 | OS1 | OS2 | OS3 |
| 1750 – 1700 | υ carbonyl C=O | 1740 | | | ↓ | | ↓ |
| ~1748 – 1740 | υ C=O of ester | 1737 | ↑← 1738 | ↓→ 1737 | | ↓→ 1731 | ←↓ 1739 |
| ~1734 | υ C=O of saturated long-chain esters or ether esters | 1728 | | ↑ | | | |
| ~1722 | υ C=O of saturated aldehydes | 1720 | ↓ | ↑ | ↑→ 1715 | ↑→ 1717 | ↑ |
| ~1715 | υ C=O of saturated aliphatic ketones | 1714 | | | | | ↑ |
| 1713 - 1709 | υ C=O of saturated aliphatic carboxylic acids | | | | | | |

(b) Important amide I and amide II bands of proteins

| Peak Frequency ($cm^{-1}$) | Assignment | Prominent adsorption peaks in control & stress experiments | | | | | |
|---|---|---|---|---|---|---|---|
| | | Ctl. | MS1 | MS2 | OS1 | OS2 | OS3 |
| 1715 – 1680 | υ-carbonyl C=O | 1696 | 1689.9 | 1691 | 1693.7 | 1695 | 1691 |
| | | 1682 | | | | ↑ | |
| ~1690 – 1610 | υC=O and υC-N of Amide I band components | 1678.7 | | | 1683 | | 1680 |
| | | 1667 | | | ↓ | | |
| | | 1657.7 | 1656.8 | 1658.3 | ↓ | 1656.6 | 1656 |
| | | 1637.6 | 1639 | | 1646 | ↓ | 1638.7 |
| | | 1621 | | | ↑ | | |
| | | 1613 | | | ↑ | | |
| | | 1599 | | | ↑ | | |
| | | 1576 | | 1578 | 1583 | | 1573 |
| | | 1570 | | | ↑ | | |
| ~1580 – 1470<br>~1577<br>~1549<br>~1538<br>~1518 - 1495 | υC-N, υ C-C, and δ N-H of Amide II band components | 1559 | | | ↑ | | |
| | | 1552.2 | ↓ | 1553.9 | | ↓ | |
| | | 1546 | 1544 | 1541 | ↓ | ↓ | ↓ |
| | | 1539 | | | | | ↑ |
| ~1520 – 1490 | Aromatic rings (Incl. Tyr and Phe) | 1537 | ↑ | | | | |
| | | 1536.6 | | | | ↑ | |
| | | 1534 | | | ↑ | | |
| | | 1517 | 1515 | – 1520 | – 1523 | 1516 | 1515 |
| | | 1505 | | | ↑ | | |

(c) Important complex infrared absorption bands of lipids, fatty acids and proteins

| Peak Frequency ($cm^{-1}$) | Assignment | Prominent adsorption peaks in control & stress experiments | | | | | |
|---|---|---|---|---|---|---|---|
| | | Ctl. | MS1 | MS2 | OS1 | OS2 | OS3 |
| | | 1468 | ↓ 1470 | ↑ | ↑ | ↓ | ↓ |
| | | 1464 | | | ↑ | | |
| ~1475 – 1370 | | 1454 | ↑ 1455 | ↓ 1455 | ↑ 1443 | ↑ 1455 | ↑ |
| ~1470<br>~1456<br>~1439 | δC-H of $CH_3$, $CH_2$<br>C-H def of >$CH_2$ | 1437.5 | ↓ 1441 | ↑ 1441 | ↓ 1440 | ↓ 1440 | ↓ 1437 |
| | | 1434 | | | ↓ | | |
| | | 1420 | | | | ↓ | |
| ~1440 – 1395 | C-O of carboxylic acid or -COO of carboxylate | 1418 | | | ↑ | | ↓ |
| | | 1414 | | ↓ | | | |
| | | 1402 | ↓ | ↓ 1398 | ↓ | | ↓ |
| | | 1393 | | | | | ↑ |
| | | 1385 | ↑ 1390 | ↑ 1386.9 | | ↓ 1392 | ↓ 1383 |
| ~1370 +/– 20 | Bending (δ, ρ, ω, τ) of OH of primary & secondary alcohols (1370 cm-1 indicate carbohydrate-rich bands ) | 1379 | | | | ↑ | |
| | | 1364.4 | ↓ 1370 | ↓ 1373 | ↓ 1358 | ↑ 1367 | ↓ 1369 |
| | | 1349 | | | | ↑ | |
| | | 1346 | ↑ | | | | |
| | | 1344 | | | ↑ | | |
| | | 1335 | ↑ 1331 | ↑ 1340 | ↑ 1331 | ↑ | ↑ 1340 |
| | | 1322 | | ↑ | | ↑ | |
| | | 1318 | | | | | ↑ |
| | | 1315 | | | ↑ | | |
| | | 1312 | | | | ↑ | |
| | | 1302 | ↑ | ↓ | | | ↑ |
| | | 1292 | | | | – 1297 | |
| ~1310 – 1240 | Amide III components of protein<br>1300 $cm^{-1}$ α-helix<br>1285 & 1245 $cm^{-1}$ random coil<br>1265 & 1235 $cm^{-1}$ β-turn spectrum | 1286.9 | ↓ 1281 | | | ↑ 1279 | ↓ 1288 |
| | | 1268 | ↑ | | ↑ | | |
| | | 1258.1 | | ↓ 1259 | | ↓ 1260 | ↑ 1264 |
| | | 1251 | | | ↑ | ↓ | |
| | | 1248 | | ↑ | | ↑ | |
| | | 1243 | ↑ | | | | |

(d) Important infrared absorption bands of carbohydrates

| Peak Frequency ($cm^{-1}$) | Assignment | Prominent adsorption peaks in control & stress experiments | | | | | |
|---|---|---|---|---|---|---|---|
| | | Ctl. | MS1 | MS2 | OS1 | OS2 | OS3 |
| 1250 – 1220 | $\upsilon_{as}$ P-O of >$PO_2^-$ phosphodiester | 1236 | | ↑ | ↑ | ↑ | ↑ |
| | | 1227.7 | 1225 | 1226 | 1221 | 1229 | 1224 |
| | | 1216 | 1213 | 1213 | | | |
| 1240 – 1171 | δ C-H of $CH_2$ coupled with C-O modes | 1193 | | 1190 | 1197 | | |
| | υ of C-O-C linkages - 1160: carbohydrate rich bands | 1183 | | | | ↑ | |
| | | 1172.7 | 1176 | 1172 | 1175 | ↓ | ↓ |
| 1200 – 1000 | $\upsilon_s$ C-O, C-C, and C-O-H, C-O-C def of carbohydrates | 1159.5 | 1163 | 1154 | 1155 | 1153 | 1166 |
| 1300 – 1200/ 1050 - 1000 | $\upsilon_{as}$ C-O of mixed (alkyl/ aryl) ethers in carbohydrates - 1240: carbohydrate rich bands | | | | | | |

(d) Important infrared absorption bands of carbohydrates (cont.)

| Peak Frequency ($cm^{-1}$) | Assignment | Prominent adsorption peaks in control & stress experiments | | | | | |
|---|---|---|---|---|---|---|---|
| | | **Ctl.** | **MS1** | **MS2** | **OS1** | **OS2** | **OS3** |
| | | 1128 | | | | ↑ | |
| | | 1124 | | ↑ | ↑ | | |
| | | 1120 | | | | | ↑ |
| 1150 – 1070 | $\upsilon_{as}$ C-O of secondary hydroxyl - 1150: carbohydrate of bacterial gum (Nichols, 1985) | 1114 | | | | | |
| | | 1107.2 | | | 1102.9 | 1109 | |
| | | 1094.5 | | 1089 | | 1091 | |
| 1090 – 1085 | $\upsilon_{as}$ P-O of $>PO_2^-$ | 1082 | 1079 | 1079 | 1085 | 1081 | ↑ |
| | | 1079 | | 1065 | 1069 | | |
| | | 1059 | 1061 | 1056 | 1053 | 1053 | |
| | | 1044 | 1046 | 1038 | 1038 | 1041 | |
| 1075 – 1000 | $\upsilon_{as}$ C-O of primary hydroxyl - of carbohydrates | 1033 | 1034 | | 1032 | 1027 | 1030 |
| | | 1027 | ↑ | | | | |
| | | 1018 | | | 1022 | | |
| | | 1006 | 1008 | 1008 | 1010 | 1009 | 1011 |

**Supplemental Table 11:** Characteristic key absorption frequencies in the mid-infrared spectral range from 1800 to 1000 $cm^{-1}$ for Sphingomonas-HTF under different types of stress. Refer to supplemental table 4 for additional information on adsorption peak assignment, and a table key.

(a) Important carbonyl (C=O) bands of lipids & fatty acids

| Peak Frequency ($cm^{-1}$) | Assignment | Prominent adsorption peaks in control & stress experiments | | | | | |
|---|---|---|---|---|---|---|---|
| | | Ctl. | MS1 | MS2 | OS1 | OS2 | OS3 |
| 1750 – 1700 | υ carbonyl C=O | 1749.2 | | ↑ | | | |
| ~1748 – 1740 | υ C=O of ester | 1747 | | | ↓ | | |
| ~1734 | υ C=O of saturated long-chain esters or ether esters | 1742 | | ↓→ 1740 | ←↑ 1748 | ←↓ 1746 | — |
| ~1722 | υ C=O of saturated aldehydes | 1738 | | | ↑ | | ↑ |
| ~1715 | υ C=O of saturated aliphatic ketones | 1728 | ↓ | — 1727 | ↓ | ↓ | ↓→ 1727 |
| 1713 - 1709 | υ C=O of saturated aliphatic carboxylic acids | 1720 | | | | | ↑ |

(b) Important amide I and amide II bands of proteins

| Peak Frequency (cm$^{-1}$) | Assignment | Prominent adsorption peaks in control & stress experiments | | | | | |
|---|---|---|---|---|---|---|---|
| | | Ctl. | MS1 | MS2 | OS1 | OS2 | OS3 |
| | | 1691 | | | | ↑ | |
| 1715 – 1680 | υ-carbonyl C=O | 1688 | ↑ | ↓ | | | |
| | | 1682 | | | | 1680 | |
| | | 1678 | | | | | ↑ |
| | | 1675 | | ↑ | | | |
| ~1690 – 1610 | υC=O and υC-N of Amide I band components | 1659 | | 1660 | 1658 | 1658 | 1658 |
| | | 1652 | ↓ | | | | |
| | | 1640 | 1639 | 1641 | 1639 | 1638 | 1641 |
| | | 1600 | | ↑ | | | |
| | | 1584 | | ↑ | | | |
| | | 1573 | | ↑ | | | |
| ~1580 – 1470<br>~1577 | | 1551 | | ↑ | | | |
| ~1549 | υC-N, υ C-C, and δ N-H of Amide II band components | 1543.3 | 1542.5 | 1539 | 1549.4 | 1549 | 1546 |
| ~1538<br>~1518 - 1495 | | 1526 | | | | | ↑ |
| | | 1516.6 | ↓ | 1522 | 1515 | ↓ | 1514 |
| ~1520 – 1490 | Aromatic rings (Incl. Tyr and Phe) | 1501 | | | | | ↑ |
| | | 1497 | | ↑ | | | |
| | | 1495 | ↑ | | | | |

(c) Important complex infrared absorption bands of lipids, fatty acids and proteins

| Peak Frequency ($cm^{-1}$) | Assignment | Prominent adsorption peaks in control & stress experiments | | | | | |
|---|---|---|---|---|---|---|---|
| | | Ctl. | MS1 | MS2 | OS1 | OS2 | OS3 |
| | | 1469 | — | ↓ | ↓ | ↓ | ↓ 1471 |
| ~1475 – 1370<br>~1470<br>~1456<br>~1439 | δC-H of $CH_3$, $CH_2$<br>C-H def of $>CH_2$ | 1456 | ↓ | 1455 | 1453 | 1454 | |
| | | 1439 | ↑ | 1438 | 1443 | 1442 | 1443 |
| | | 1430 | | | | | ↑ |
| ~1440 – 1395 | C-O of carboxylic acid or<br>-COO of carboxylate | 1424 | | ↑ | ↑ | | |
| | | 1413 | ↓ | 1409 | ↓ | — | 1407 |
| | | 1400 | | | ↑ | | |
| | | 1393 | ↑ | 1397 | 1389.8 | 1395 | |
| | | 1383 | ↑ | ↑ | | 1387 | |
| ~1370 +/– 20 | Bending (δ, ρ, ω, τ) of OH of primary & secondary alcohols (1370 cm-1 indicate carbohydrate-rich bands ) | 1370 | 1369 | ↑ | ↓ | 1375 | 1376 |
| | | 1361 | ↑ | | | | |
| | | 1357 | | ↑ | | | |
| | | 1351 | | | 1347 | | 1357 |
| | | 1337 | ↓ | 1332 | ↓ | 1336 | 1345 |
| | | 1325 | ↑ | ↑ | ↓ | ↑ | |
| | | 1319 | | | | | ↑ |
| | | 1307 | ↑ | 1302 | 1306 | 1312 | ↓ |
| | | 1299 | | | | ↑ | |
| ~1310 – 1240 | Amide III components of protein<br>1300 $cm^{-1}$ α-helix<br>1285 & 1245 $cm^{-1}$ random coil<br>1265 & 1235 $cm^{-1}$ β-turn spectrum | 1288 | ↑ | 1294 | 1297 | 1284 | 1295 |
| | | 1275 | | | | | ↑ |
| | | 1268 | | ↑ | ↑ | | |
| | | 1264 | ↑ | | | | |
| | | 1261 | | | | | ↑ |
| | | 1258 | | | ↑ | | |
| | | 1249 | ↑ | 1255 | — 1246 | 1255 | 1252 |

(d) Important infrared absorption bands of carbohydrates

| Peak Frequency (cm$^{-1}$) | Assignment | Prominent adsorption peaks in control & stress experiments | | | | | |
|---|---|---|---|---|---|---|---|
| | | Ctl. | MS1 | MS2 | OS1 | OS2 | OS3 |
| 1250 – 1220 | $\upsilon_{as}$ P-O of >$PO_2^-$ phosphodiester | 1235 | ↓ | ↓ 1244 | | ↓ 1244 | ↑ 1236 |
| | | 1225 | ↓ | ↓ 1233 | ↓ | ↓ 1228 | ↑ 1227 |
| 1240 – 1171 | δ C-H of $CH_2$ coupled with C-O modes | 1212 | ↑ 1213 | ↑ 1210 | | | ↓ 1215 |
| | υ of C-O-C linkages - 1160: carbohydrate rich bands (Nichols, 1985) | 1194 | ↑ 1192 | ↑ 1198 | | | ↑ 1197 |
| | | 1184 | | | | | ↑ |
| | | 1171 | — | ↑ | | | |
| | | 1160 | ↑ | | ↓ 1166 | ↓ 1166 | ↑ 1169 |
| | | 1151 | ↓ | ↑ 1148 | ↑ 1154 | ↓ 1155 | |
| | | 1137 | | | | | ↑ |
| | | 1134 | | | ↑ | | |
| 1150 – 1070 | $\upsilon_{as}$ C-O of secondary hydroxyl - 1150: carbohydrate of bacterial gum | 1122 | ↑ 1123 | | ↑ | ↑ 1126 | ↑ 1123 |
| | | 1114 | | ↑ | | | |
| 1140 – 1070 | $\upsilon_{as}$ C-O of saturated unbranched ethers in carbohydrates - | 1107 | ↑ | | | | ↓ 1104 |
| | | 1101 | | ↑ | | | |
| 1090 – 1085 | $\upsilon_{as}$ P-O of >$PO_2^-$ | 1088.5 | ↓ | | | | ↑ 1091 |
| | | 1081 | ↓ | ↑ 1083 | ↑ 1084 | ↑ 1082 | |
| | | 1076 | | | | | ↑ |
| | | 1070 | ↑ 1071 | | | | |
| 1075 – 1000 | $\upsilon_{as}$ C-O of primary hydroxyl - of carbohydrates | 1060 | ↑ 1061 | ↑ 1063 | | | ↑ 1057 |
| | | 1056 | | ↑ | | | |
| | | 1040 | ↑ | ↑ | | | ↑ 1043 |
| 1200 – 1000 | $\upsilon_s$ C-O, C-C, and C-O-H, C-O-C def of carbohydrates | 1030 | ↓ | ↑ 1032 | ↑ | ↑ 1032 | |
| | | 1020 | ↓ | | | ↓ 1024 | ↑ 1024 |
| 1300 – 1200/ 1050 - 1000 | $\upsilon_{as}$ C-O of mixed (alkyl/ aryl) ethers in carbohydrates - 1240: carbohydrate rich bands | 1015 | | ↑ | | | |
| | | 1005 | | | | | ↑ |
